# Architecture-Dependent Stability, Cellular Uptake, and Redox Modulation of Poly(p-Coumaric Acid) Hybrid Nanoparticles for Ovarian Carcinoma Intervention

**DOI:** 10.64898/2026.04.12.717943

**Authors:** M. Megahed, A. Harun, K. Herrera, Md. H. Rashid, R. Posey, I. De Leon, J. Tropp, I. Srivastava

## Abstract

The clinical efficacy of fluorescence-guided surgery (FGS) is often compromised by the poor photostability and biologically inert nature of conventional contrast agents such as Indocyanine Green (ICG). While nanocarriers can enhance dye stability, they often function primarily as passive delivery vehicles, requiring additive complexity to achieve therapeutic effects. Here, we report a structure-guided approach to develop self-theranostic hybrid nanoparticles where the polycondensation kinetics of the polymer core, poly(p-coumaric acid) (PCA), serve as a critical design parameter governing nanoparticle assembly and downstream optical and biological performance. By systematically varying the reaction duration, we synthesized PCA variants with distinct polymer growth profiles that influence nanoparticle morphology, ICG encapsulation, and fluorescence stability. The optimized PCA1.5h formulation significantly improved the stability of encapsulated ICG, maintaining robust NIR-I fluorescence under storage and surgical illumination conditions. Beyond acting as a structural scaffold, the PCA matrix retained intrinsic redox-modulating activity, leading to increased reactive oxygen species (ROS) signal generation and reduced viability in multiple ovarian cancer cells. The imaging performance of these nanoparticles was further evaluated using 3D bioprinted intraperitoneal tumor phantoms designed to simulate key optical and spatial features relevant to fluorescence-guided imaging. This work establishes reaction-time-dependent PCA growth profiles as an important formulation parameter for integrating imaging performance and intrinsic biological activity within a simplified hybrid nanomaterial platform.

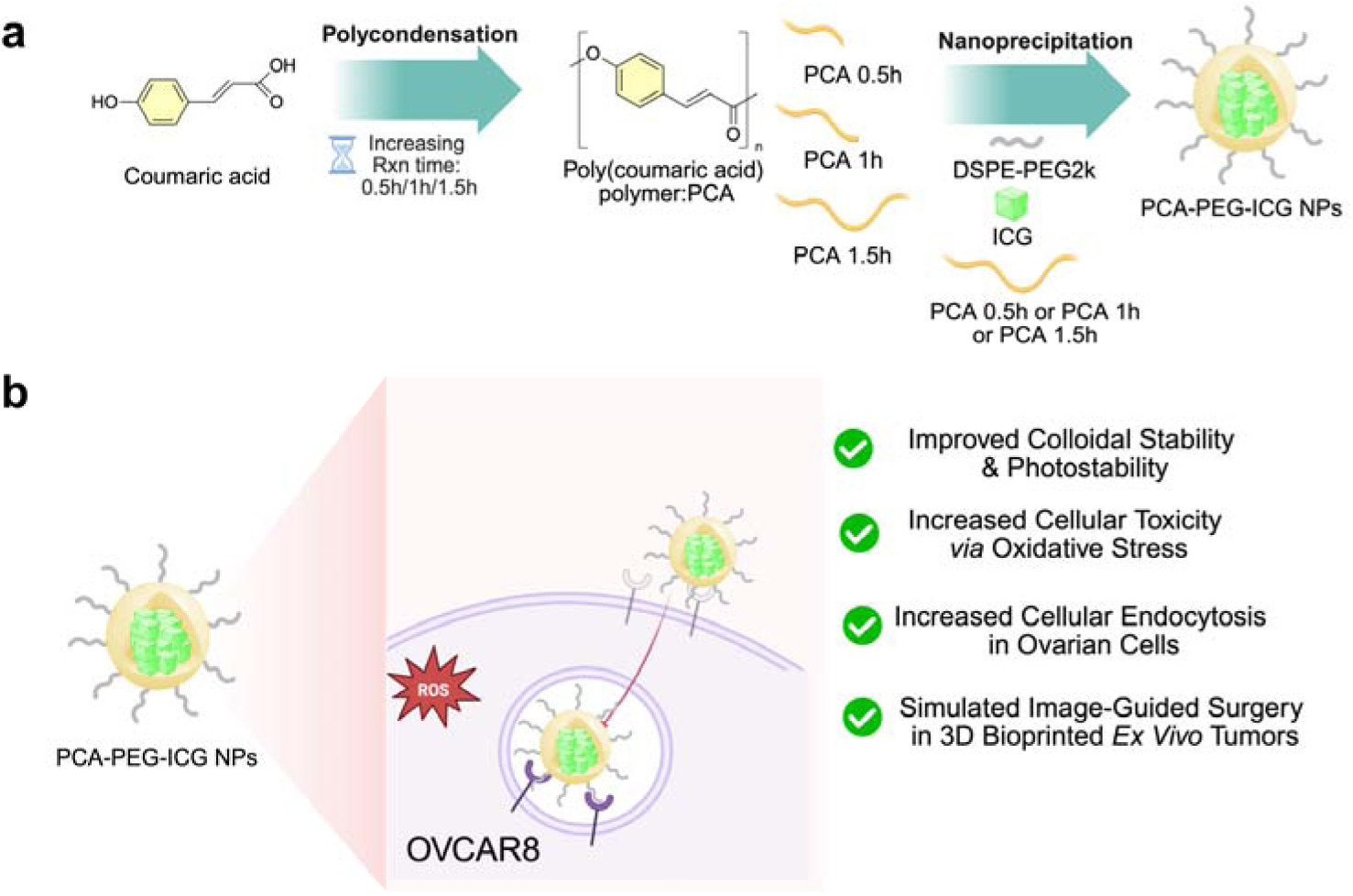
TOC.

## INTRODUCTION

Fluorescence-guided surgery (FGS) has emerged as a transformative modality for enhancing surgical precision, enabling real-time, high-contrast visualization of malignant margins.^1, 2^ Among clinically approved fluorophores, Indocyanine Green (ICG) remains the gold standard due to its near-infrared (NIR-I) emission and deep tissue penetration.^3–5^ However, the translational utility of free ICG is severely hampered by rapid photobleaching, concentration-dependent quenching, and poor aqueous stability.^4, 6^ These limitations lead to premature signal decay during prolonged cytoreductive procedures, often necessitating repeated dosing or compromising the detection of residual micro-tumors.^7–9^

To mitigate these drawbacks, diverse nanocarrier platforms have been engineered to encapsulate and stabilize ICG.^9, 10^ While successful in improving optical longevity, most existing systems are designed as biologically inert “shipping containers” that contribute no therapeutic value.^9, 11, 12^ This stark dichotomy between imaging functionality and therapeutic efficacy represents a missed opportunity in nanomedicine design, where multifunctionality is typically achieved through “additive complexity", the conjugation of multiple disparate components, rather than the intrinsic material properties of the scaffold itself.^13–15^

An elegant solution lies in the development of self-therapeutic polymers, where the macromolecular backbone possesses innate biological activity.^16, 17^ Bio-derived phenolic polymers are particularly compelling candidates due to their inherent redox activity and ability to modulate intracellular oxidative stress.^18,19^ Poly(p-coumaric acid) (PCA), synthesized via the polycondensation of naturally occurring p-coumaric acid, represents a unique class of “active” scaffolds.^16, 20^ Despite the potential of such systems, a fundamental gap remains: the role of macromolecular architecture in governing the transition from a chemical scaffold to a functional nanomedicine. In current literature, polymer structure is frequently treated as a fixed attribute rather than a tunable design variable.^16, 21^ Consequently, the correlation between polymerization kinetics, nanoparticle assembly, and emergent theranostic performance remains largely unexplored.^14, 22^

Here, we demonstrate that the polycondensation reaction time of PCA represents a key design parameter influencing nanoparticle performance in ovarian carcinoma models. By systematically varying reaction duration, we synthesized PCA variants with distinct polymer growth profiles that affected hybrid lipid–polymer nanoparticle assembly. The optimized PCA1.5h formulation improved ICG encapsulation, dye retention, and fluorescence stability under storage and surgical illumination conditions. In cell-based assays, PCA1.5h-derived nanoparticles exhibited increased cell-associated fluorescence, reduced cell viability, and elevated reactive oxygen species (ROS) signal relative to lower reaction-time formulations. Finally, we evaluated these nanoparticles using controlled tumor-mimicking phantoms and 3D-bioprinted intraperitoneal tumor models designed to simulate key optical and spatial features relevant to fluorescence-guided imaging. Together, these findings establish reaction-time-dependent PCA growth profiles as a critical design parameter for tuning nanoparticle assembly, optical stability, and biological activity.

## MATERIALS AND METHODS

### Materials

All materials were purchased from Sigma-Aldrich and used without modification, unless otherwise stated. Porcine skin tissue was obtained from local grocery stores. Preserved bovine brains with dura mater, porcine ovaries, and dry-preserved porcine lung tissue were purchased from Nasco Education. This study did not involve human participants, human clinical specimens, or animal experiments. Experiments using established cell lines did not require human subjects’ approval.

### Preparation and Characterization of Poly-Coumaric acid (PCA)

Poly coumaric acid was synthesized using polycondensation reaction at three different reaction times. Under 4°C ice bath conditions, 10 ml of anhydrous pyridine was stirred for 5 min in a round bottom flask, followed by the addition of 5 mmol of thionyl chloride. After 15 min the round bottom flask was removed from the ice bath, and 2.5 mmol of p-coumaric acid was added and left to stir at room temperature for 30 min, 1 hour and 1.5 hours (referred as PCA0.5h, PCA1h, and PCA1.5h respectively). 1% HCL was then added to stop the reaction, and the samples were washed twice (13000 rpm, 30 mins) using Ultrapure water and then lyophilized into powders. 1H-NMR spectroscopy was used to characterize the polymers’ structures and composition using a JEOL ECS 400 MHz spectrometer. Ultraviolet-visible (UV-VIS) absorption was recorded using the Genesys 30 Visible Spectrophotometer (ThermoFisher Scientific). Number-average molecular weight (*M*_n_), weight-average molecular weight (*M*_w_), and dispersity (Đ) were determined by gel permeation chromatography (GPC) on a Dionex liquid chromatography system equipped with a Thermo Fisher variable-wavelength detector (VWD) monitoring at 300 nm to match the peak absorption of the PCA samples. Separations were carried out at a column oven temperature of 50°C using two Varian Polypore columns (300 × 7.5 mm) connected in series, with N,N-dimethylformamide (DMF) containing 0.4% lithium chloride (LiCl) as the mobile phase at 1.0 mL min^-1^. The system was calibrated using six narrow-dispersity polystyrene standards ranging from *M*_w_ = 2.56 × 10^3^ – 1.10 × 10^5^ (TSKgel Standard Kit, Tosoh). Samples were dissolved in the DMF eluent at 1.0 mg mL^-1^ and filtered through a 0.45 **μ**m PTFE membrane prior to injection.

### Preparation and Characterization of Poly-Coumaric Acid Nanoparticles (PCA-PEG NPs)

Following the synthesis of poly-coumaric acid (PCA) at reaction times of 0.5 h, 1 h, and 1.5 h, the polymers were used to formulate NPs using DSPE-PEG *via* nanoprecipitation ^23–25^. The aqueous phase was identical for all reaction times and consisted of 8 mL of PBS 1X. The organic phase of the PCA-PEG NPs consisted of 1 mL of the PCA stock solution (1 mg/mL in DMSO), 1 mL of DMSO and 500 **μ**L of the DSPE-PEG stock solution (5 mg/mL in DMSO). For PCA-PEG-ICG NPs, the organic phase consisted of 1 mL of the PCA stock solution, 500 **μ**L of the DSPE-PEG stock solution, 500 **μ**L of DMSO, and 200 **μ**L of ICG stock solution (1 mM in DMSO). The organic phase was added dropwise to the aqueous phase while sonicating. Upon completion, the NPs were ultracentrifuged at 4,500 rpm for 10 min and washed four times with PBS 1X to remove unreacted precursors and free ICG. The washed NPs were then filtered through a 0.22-**μ**m PVDF filter and stored at 4 °C.

### Optimization of DSPE-PEG Concentration in PCA-PEG NPs

To evaluate the effect of DSPE-PEG content on NPs physicochemical properties, various concentrations of DSPE-PEG namely 5, 10 and 20mg/mL were used to formulate the NPs using nanoprecipitation while using the same polymer PCA 1.5h. Briefly, the organic phase consisted of 1 mL PCA1.5h stock solution (1 mg/mL in DMSO), 1 mL DMSO, and 500**μ**l of DSPE-PEG stock solution at concentrations of 5, 10, or 20 mg/mL, while the aqueous phase was of 8 mL PBS (1X). The organic phase was added dropwise into the aqueous phase under continuous sonication. Upon completion, the NPs were ultracentrifuged at 4,500 rpm for 10 min and washed four times with PBS 1X to remove unreacted precursors. The washed NPs were then filtered through a 0.22-**μ**m PVDF filter and stored at 4 °C.

For clarity, formulations prepared during PEG optimization are denoted as PEG5, PEG10, and PEG20, corresponding to PEG concentrations of 5, 10, and 20 mg/mL, respectively. Following optimization, 5 mg/mL PEG was found to be the most ideal candidate selected for remainder of the experiments, and subsequent optimized formulations are referred to as PCA-PEG or PCA-PEG5 unless otherwise specified.

### Physicochemical and Optical Characterization of PCA-PEG and PCA-PEG-ICG NPs

Hydrodynamic diameter (HD) and polydispersity index (PDI) were determined via dynamic light scattering (DLS), using the Litesizer DLS 700 Particle Analyzer instrument (Anton Paar). **ζ**-potential measurements also used the Litesizer DLS 700 Particle Analyzer. Ultraviolet-visible (UV-VIS) absorption was recorded using the Genesys 30 Visible Spectrophotometer (ThermoFisher Scientific). All absorbance measurements were recorded with scanning intervals of 2 nm from 250 to 1100 nm. Fluorescence characterization of PCA-PEG-ICG NPs was performed using a fluorescence spectrophotometer RF6000 (Shimadzu) at an excitation wavelength of 785 nm and an emission range of 800–900 nm.

### Transmission Electron Microscopy Imaging (TEM)

Carbon coated copper grids of 200 mesh were glow discharged immediately prior to use to render the surface hydrophilic. A drop of PCA NP was then applied to the grid surface, and it was incubated for 60 seconds. Excess liquid was removed by blotting with filter paper. Negative staining was performed by adding a drop of freshly prepared 1% w/v uranyl acetate solution to the grid surface, followed by a 30-second incubation. Excess stain was then carefully blotted away, and the grids were dried under vacuum at room temperature overnight. All samples were imaged using a Hitachi H-7650 transmission electron microscope at 100 kV.

### PCA-PEG NPs Stability Assessment

The colloidal stability of the PCA1.5-PEG, PCA1.5-PEG-ICG, PCA1-PEG-ICG and PCA0.5-PEG-ICG NPs was evaluated by incubating them with PBS buffer 1x (pH 7.4) at 4°C and 37°C and with 1% plasma at 37°C. The size stability was tested by DLS over 7 days.

### Release Kinetics of ICG *in vitro*

ICG release from PCA1.5-PEG-ICG NPs was evaluated using a dialysis method under physiological and acidic pH conditions. Briefly, PCA1.5-PEG-ICG NP suspension (5.5 mL) was loaded into dialysis cassettes with a molecular weight cutoff of 20 kDa MWCO and immersed in PBS (pH 7.4) or PBS adjusted to pH 5.0. Samples were incubated for 24 h at room temperature under continuous agitation at 950 rpm and protected from light. Retentate samples containing PCA1.5-PEG-ICG NPs were collected at 0 and 24 h. Dialysate samples were collected at 0, 1, 2, 4, 8, and 24 h to assess released ICG. UV–Vis absorbance spectra were recorded from 650–900 nm, and fluorescence emission spectra were collected using 785 nm excitation and an emission range of 800–900 nm. At each time point, 1 mL of dialysate was collected and replaced with an equal volume of fresh buffer at the corresponding pH to maintain sink conditions. ICG concentration in the dialysate was calculated using an ICG calibration curve, and release was reported as the amount of ICG detected in the dialysate over 24 h.

### *In Vitro* Cytotoxicity Evaluation

The cytotoxic effects of PCA0.5-PEG, PCA1-PEG and PCA1.5-PEG were assessed using the colorimetric MTT assay. OVCAR8 cells, OVCAR3 Cells and HUVECs, (5 × 10^4^ cells/well) were seeded in 96-well plates in 200 **μ**L of reconstituted DMEM medium for 24 h. After 48 hours, OVCAR8 cells were incubated with NPs at increasing concentrations (1, 5, 10, 25, 50, and 100 µg/mL), whereas OVCAR3 cells and HUVECs were incubated with the NPs at the same increasing concentration for 24 hours. Subsequently, cells were incubated with 10 **μ**L of MTT stock solution in PBS (5 mg/mL) per well for 4 h, followed by dissolving the formazan crystals with 200 µL of DMSO. The absorbance was measured using the BioTek Epoch 2 Microplate Spectrophotometer (Agilent) at 592 nm.

### Cellular Uptake Kinetics of Optimized PCA1.5-PEG-ICG NPs

As PCA1.5-PEG-ICG NPs showed an optimized architectural and stability design, their cellular internalization was first assessed. OVCAR8 cancer cells were seeded in 6-well plates at a density of 2× 10**O** cells per well and cultured in DMEM supplemented with 10% fetal bovine serum (FBS). After 24 hours, the cells were incubated with PCA1.5-PEG-ICG NPs for 4, 8, 12and 24 h at an ICG concentration of 7.5 **μ**M. Following incubation, cells were collected into microcentrifuge tubes, and the fluorescence emission images were then measured using an IVIS imaging system at an excitation filter of 745 nm and an emission filter of 840 nm. To further verify intracellular localization, OVCAR8 cells seeded in 24-well plates (1 × 10**O** cells/well). After 24 hours, the cells were treated with PCA1.5-PEG-ICG NPs for 4, 8, 12and 24 h (7.5 **μ**M of ICG). After NPs removal, Hoechst dye (5**μ**g/ml) in PBS 1X was added for 10 minutes. The dye was removed and the cells were washed twice with PBS1X followed by fluorescence imaging and quantification using the EVOS M5000 Fluorescence microscope (ThermoFisher).

### Formulation-Dependent Cellular Uptake of PCA-PEG-ICG NPs at 24h

OVCAR8 cancer cells were seeded in 6-well plates at a density of 2× 10**O** cells per well and cultured in DMEM supplemented with 10% fetal bovine serum (FBS). As the optimal cellular intake was observed at 24 h with using PCA1.5-PEG-ICG NPs, the cells were incubated with PCA0.5-PEG-ICG and PCA1-PEG-ICG NPs for 24 h at an ICG concentration of 7.5 **μΜ**. Following incubation, cells were collected into microcentrifuge tubes, and the fluorescence emission images were then measured using an IVIS imaging system at an excitation filter of 745 nm and an emission filter of 840 nm. For Fluorescence microscopy imaging, OVCAR8 cells were seeded in 24-well plates (1 × 10**O** cells/well). After 24 hours, the cells were treated with the NPs for 24 h (7.5 **μ**M of ICG). After removal of NPs, Hoechst dye (5**μ**g/ml) in PBS 1X was added for 10 minutes. The dye was removed and the cells were washed twice with PBS1X followed by fluorescence imaging and quantification using the EVOS M5000 Fluorescence microscope (ThermoFisher Scientific).

### *In Vitro* Assessment of ROS production post PCA-PEG NPs treatment

Intracellular production of ROS was assayed using 2**′**,7**′**-dichlorodihydrofluorescein diacetate (H_2_DCFDA) probe. OVCAR8 cells were seeded in a 24 well plate at density of 1 x 10^5^ cells/well for 24 hours and then cultured with 50 and 100 **μ**g/ml of PCA1.5-PEG NPs for 2,4 and 8 h. To further evaluate ROS generation in a second ovarian cancer cell line, OVCAR3 cells were seeded in a 24 well plate at density of 1 x 10^5^ cells per well for 24 hours and then treated with 50 **μ**g/ml of PCA1.5-PEG NPs for 8 h. Subsequently, the cells were incubated with 20 **μ**M H_2_DCFDA for 30 min at 37°C and washed twice using PBS1x. Finally, the cells were imaged and quantified using the EVOS M5000 Fluorescence microscope (ThermoFisher Scientific).

### Encapsulation Efficiency of PCA-PEG-ICG NPs

The encapsulation efficiency (EE) of ICG using PCA1.5-PEG-ICG, PCA1-PEG-ICG and PCA0.5-PEG-ICG was assessed using UV–Vis spectroscopy using the Genesys 30 Visible Spectrophotometer (ThermoFisher Scientific) at an absorption of 702 nm. A calibration curve was prepared using ICG diluted with DMSO at increasing concentrations (0.1-100 **μ**M). The absorption spectrum of the NPs was then collected under the same conditions and ICG content was then quantified based on the calibration curve.

The encapsulation efficiency was calculated according to the following equation:

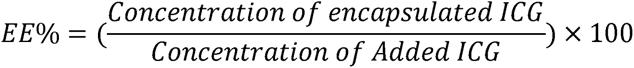

### Tumor Mimicking Phantom Stock Solution Preparation

A tumor mimicking phantom stock solution was made by combining 17 mL of PBS 1X, 900 µL of intralipid 20%, 150 mg of porcine hemoglobin, and 720 mg of type-A gelatin. The gelatin was activated following a continuous stirring at 65°C and 950 rpm for at least 30 minutes until the gelatin is fully dissolved in PBS1X. After activation, intralipid and hemoglobin were added to the solution and the mixture was inverted and vortexed, it was further mixed for 10 minutes at 950 rpm at 65°C. The stock was further used to synthesize different compositions of phantom with PCA-PEG NPs and PCA-ICG-PEG NPs.

### Quantitative Fluorescence Imaging using PCA-PEG-ICG Tumor-Mimicking Phantoms

1.5mL phantom stock solution was added with 0.5mL PCA1.5-PEG-ICG NP using a two-fold serial dilution of the NP with an initial concentration of 10 µM in silicon mold wells. The phantoms were solidified in the freezer at -20°C overnight. A control phantom was made using 2mL of phantom stock solution only in the silicon mold well following solidification in the freezer. The IVIS image of the phantoms were collected placing them in a well plate using the IVIS imaging system at the excitation wavelength of 745 nm and emission wavelength of 840 nm.

### *Ex Vivo* Porcine Tissue Penetration Profiling

Three different porcine tissue samples were used to simulate the characteristics of human skin, muscle, and fat tissues. A piece of pork belly was used to replicate skin tissue, pork tenderloin was used to resemble muscle tissues, and a fatty pork chop was used as fat tissue. Using a deli meat slicer in the lab, all tissues were consistently sliced to a thickness of 2 mm after being obtained from nearby supermarkets. The tissue slices were stacked in accordance with the variation in tumor depth in a well plate covering the PCA1.5-PEG-ICG NP phantom made with 1.5mL of phantom stock solution and 0.5mL of PCA1.5-PEG-ICG NP solution. The same tissue stacking simulation was performed in an ICG phantom made with 1.5mL of phantom stock solution and 0.5mL of ICG solution of 10 µM.

From surface-level to deep tumor lesions, tissues were stacked at different depths from 0 mm to 4 mm where cancers can develop. Fluorescence images were acquired using an IVIS imager at 745 nm excitation and 840 nm emission filter.

### Photobleaching Assessment of PCA-ICG NP

PCA1.5hr-PEG-ICG NP TMP, ICG TMP and Control TMPs were made following the methods described above. The TMPs were placed in a well plate and exposed to a 1000 lumen white light source for different time periods (1min, 0.5h, 2h, 4h). Fluorescence images were collected at each time point after being exposed to the white light source using an IVIS imager at 745 nm excitation and 840 nm emission window.

### 3D Bioprinting of Intraperitoneal Tumors

To create the intraperitoneal tumors, skin mimicking phantoms were printed, and TMPs with nanoparticles were printed on top of ‘skin’ mimics. The skin mimics were printed using 500 µL of melanin solution (2mg/mL) mixed with 2 ml of high concentration gelatin solution (6%). The mixture was placed in a syringe and added into the printer. By setting specific printing parameters and freezing the mixture to 2°C the skin mimicking phantom was printed. To print the small Tumor Mimicking Phantoms on top of the ‘skin’ mimic, the printing parameters were kept similar except the dimensioning had been decreased to 1mmx1mm to 0.1mmx0.1mm. 1.5 mL of the Tumor Mimicking Phantom solution was mixed with 500 µL of PCA1.5-PEG-ICG NPs, then added to a syringe and printed. By changing the translations on the x, y, and z axis of the printer, separate small tumor phantoms at multiple desired locations were printed on top of the skin print.

### Hemocompatibility assay of PCA1.5-PEG NPs

The hemocompatibility of PCA1.5-PEG NPs was evaluated using a hemolysis assay with Porcine blood (RBCs). 1 ml of NPs dispersed in PBS at concentrations of 1, 5, 10, 50 and 100 **μ**g/mL were incubated with 50 **μ**L RBCs for 1 h at 37 °C. PBS and deionized water served as the negative and positive controls, respectively. Following the incubation, the microcentrifuge tubes were centrifuged at 3000 rpm for 20 minutes and the absorbance of the supernatant was measured at 540 nm using a microplate reader to quantify released hemoglobin. Finally, the supernatant was imaged using an inverted microscope (LaxcoTM), to image the integrity of the RBCs.

### Statistical Analysis

Data are presented as mean ± standard deviation unless otherwise stated. The number of independent experiments, printed constructs, region-of-interest (ROI) measurements, or technical measurements is indicated in the corresponding figure legends. Statistical analyses were performed using GraphPad Prism 10. For comparisons between two groups, an unpaired two-tailed Student’s t-test was used when variances were comparable, while Welch’s t-test was used when unequal variance was expected or observed. For paired measurements collected from the same samples over time, a paired two-tailed t-test was used. For comparisons involving more than two groups or multiple treatment conditions, one-way or two-way analysis of variance (ANOVA) was used, followed by the appropriate post-hoc multiple-comparisons test. Dunnett’s post-hoc test was used for comparisons against a single control group, Tukey’s post-hoc test was used for all pairwise comparisons, and Šídák’s post-hoc test was used for selected pairwise comparisons across treatment or time conditions, as appropriate. For the 3D-bioprinted intraperitoneal tumor model, tumor-to-healthy tissue fluorescence ratios were calculated from ROI measurements of tumor-mimicking and adjacent healthy/background regions. Comparisons between IP #1 and IP #2 were performed using an unpaired two-tailed Welch’s t-test because the constructs were independently printed using the same composition and printing parameters, and ROI-level measurements could have unequal variance. Statistical significance was defined as p < 0.05, with significance indicated as *p < 0.05, **p < 0.01, ***p < 0.001, and ****p < 0.0001. Non-significant comparisons are indicated as ns.

## RESULTS AND DISCUSSION

### Synthesis and Characterization of Poly(Coumaric Acid) at Different Reaction Times

PCA was synthesized through a one-step polycondensation reaction of CA, using thionyl chloride as an activating agent in pyridine. Pyridine acted as both the solvent and base to facilitate the polycondensation reaction.^17, 19, 26^ We aimed to demonstrate that the duration of the polycondensation reaction influences polymeric nature of PCA and, consequently, its physicochemical and antioxidant properties after nanoparticles (NPs) formation **(Figure 2a)**. UV-Vis absorbance spectroscopy was used to evaluate the structural changes associated with PCA polymerization at different reaction times. As shown in **Figure 2 b,c**, the spectra of free CA and PCA samples were compared. Free CA displayed a maximum absorbance peak at 314 nm, whereas PCA 0.5h, PCA1h, and PCA1.5h showed absorbance maxima at 316, 318, and 320 nm, respectively. The progressive red shift observed with increasing reaction time suggests an increase in conjugation length, indicative of a greater extent of polymerization. Notably, PCA1.5h exhibited the most pronounced shift, consistent with more extended **π**-conjugation along the polymer backbone.^27^ ¹H-NMR spectroscopy further confirmed successful polymer formation. Free CA showed a characteristic downfield singlet for the carboxylic acid proton (-COOH) at ∼10-12 ppm and a broad phenolic OH signal at ∼9-10 ppm. In contrast, the disappearance of the carboxylic acid signal PCA **(Figure 2 d,e and Figure S2-S5)** indicates that these groups participated in the polycondensation reaction, forming ester linkages along the backbone **(Figure S1)**. ^16, 17, 20^ Additionally, spectra in **Figure S3-S5** show progressively sharper signals corresponding to the alkene protons, suggesting increased structural regularity and a more defined polymer backbone with longer reaction times.

**Figure 1.**
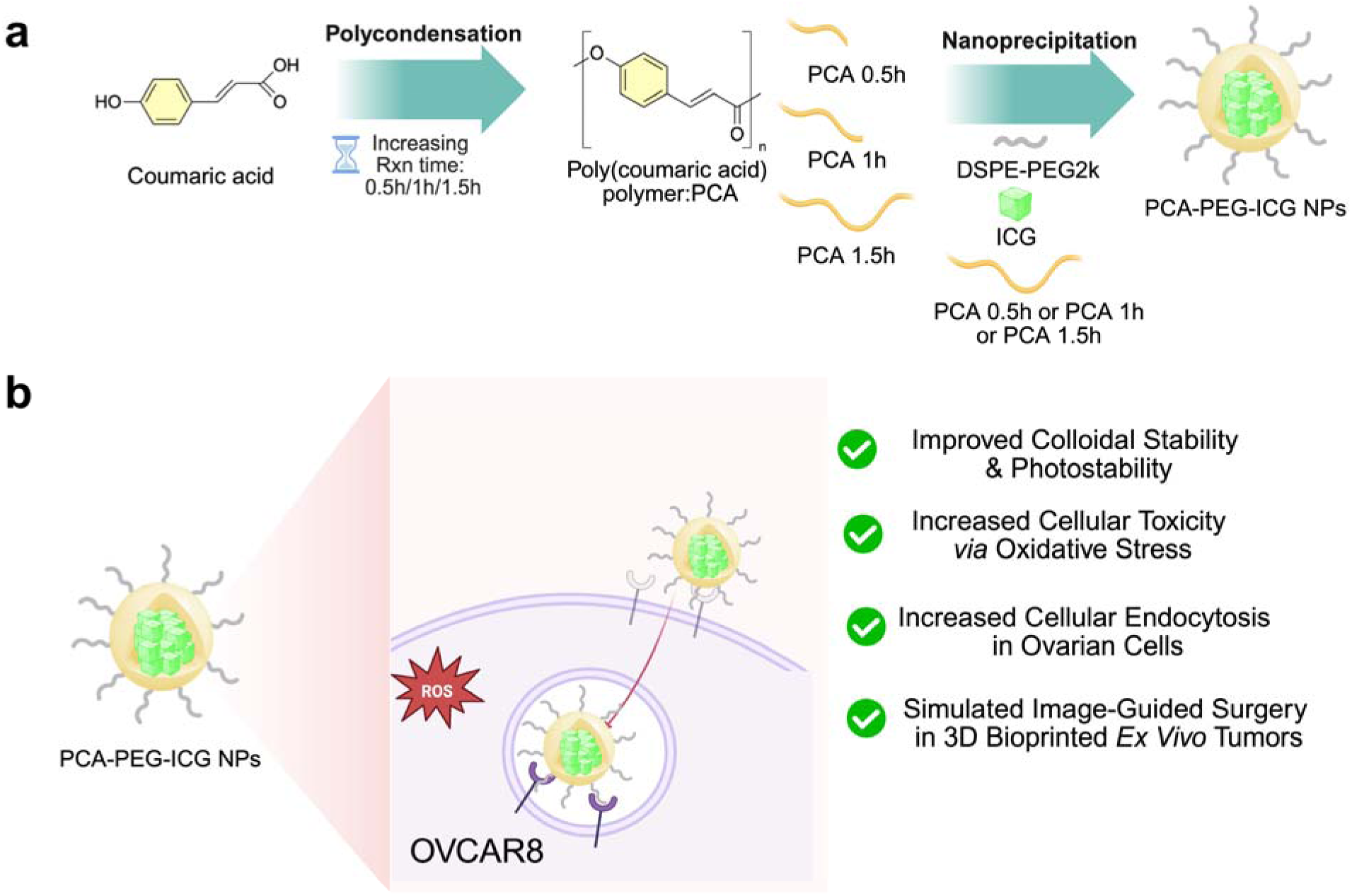
Schematic illustration of the synthesis, formulation, and biological evaluation of PCA-PEG-ICG nanoparticles. (a) Coumaric acid undergoes polycondensation to form poly(coumaric acid) (PCA) with varying reaction times (0.5, 1, and 1.5 h). The resulting PCA polymers are then formulated into nanoparticles via nanoprecipitation with DSPE-PEG2k and ICG, yielding PCA-PEG-ICG NPs. (b) The nanoparticles exhibit increased cellular uptake via endocytosis, and elevated cytotoxicity mediated by oxidative stress in OVCAR8 cells. Additionally, the NPs showed enhanced colloidal stability and photostability and are evaluated in *ex vivo* bioprinted tumor models to simulate image-guided surgery (IGS) applications.

**Figure 2.**
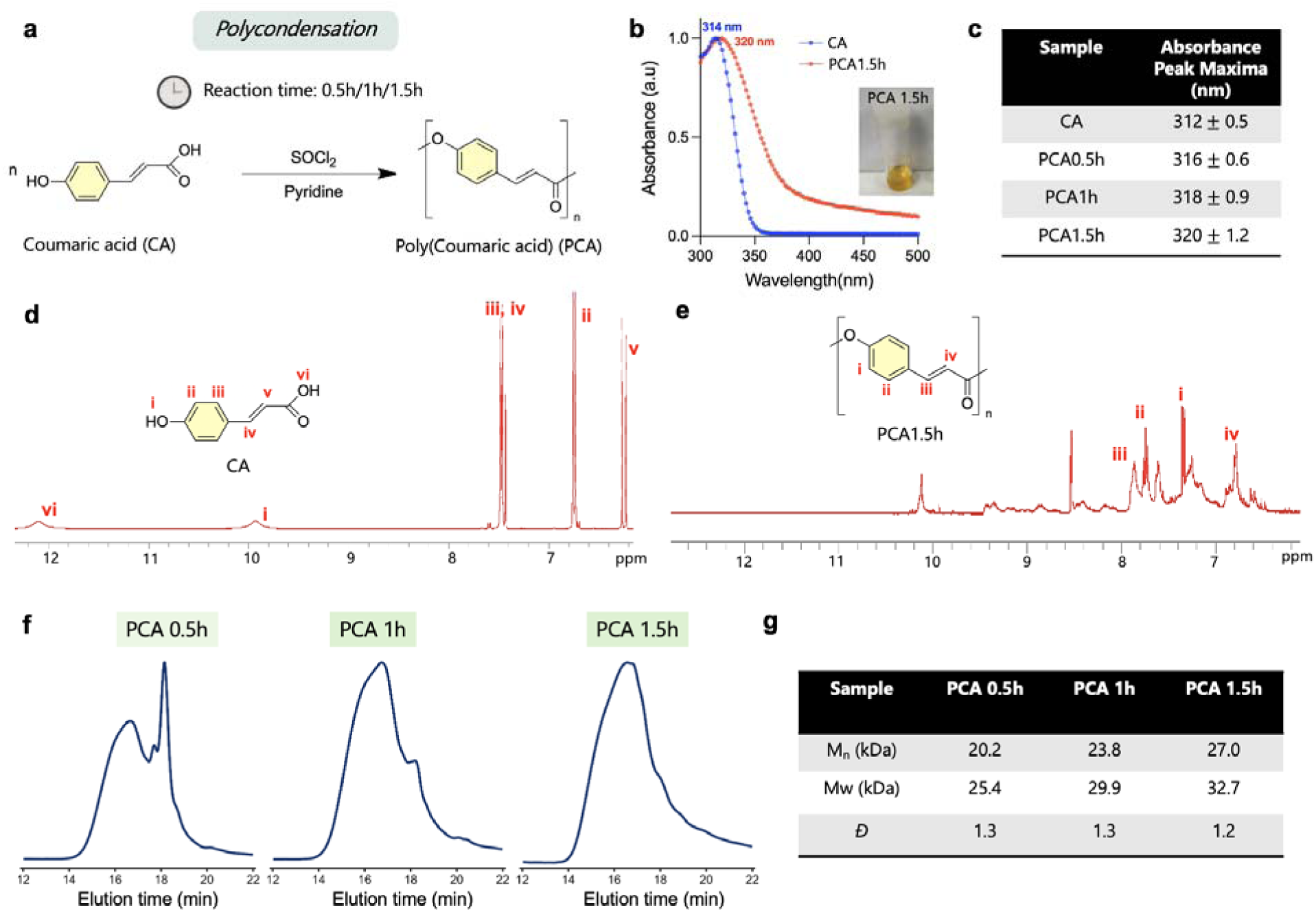
Time-dependent polycondensation reaction to synthesize poly(coumaric acid) and their characterization. (a) Schematic overview of poly-coumaric acid (PCA) synthesis through one step poly condensation using coumaric acid (CA) (b) UV-Vis absorbance spectra of coumaric acid (CA) and PCA 1.5h polymer (c) Peak UV-Vis absorbance maxima of CA and PCA at various reaction times (d, e) 1H-NMR spectroscopy of CA and PCA1.5h showing the disappearance of the carboxylic group peak at 12 ppm as the polymer is formed. (f) Gel permeation chromatography/size-exclusion chromatography elution profiles of PCA synthesized at different reaction times. (g) Summary of number-average molecular weight (M**O**), weight-average molecular weight (M_w_), and dispersity (Đ) of PCA synthesized at different reaction times.

Gel permeation chromatography (GPC) was used to characterize the molecular weight distribution of the PCA samples and monitor molecular-weight evolution as a function of polycondensation time. As PCA is formed through step-growth polycondensation of p-coumaric acid, the resulting molecular weight distributions are expected to approximate Flory–Schulz statistics. For such step-growth materials, GPC provides a direct assessment of molecular weight distribution and associated parameters (*M*_n_, *M*_w_, Đ), whereas number-average molecular weights obtained from end group analysis become increasingly unreliable at higher degrees of polymerization due to the diminished contribution of end-group signals (e.g., NMR, MALDI). As the Carothers equation directly relates degree of polymerization (DP) with monomer conversion,^28^ reaction time was used as a kinetic handle to afford polymer batches with systematically increasing molecular weight. GPC analysis supports reaction-time-dependence on molecular weight of PCA (**Figure 2f, g**). The M_w_ of PCA 0.5h, 1.0h, and 1.5h increased monotonically from 25.4, 29.9, and 32.7 kDa, respectively; these are the highest molecular weights reported to date for such poly(acid) materials.^16,20,21,29^ At 0.5 h the distribution is bimodal, with a prominent low-molar-mass population eluting near 18 min, whereas at 1 and 1.5 h this oligomeric shoulder is progressively consumed and the traces converge onto a single, higher-molar-mass population. The convergence of the 1 h and 1.5 h chromatograms indicates that the molecular-weight distribution reaches a stable endpoint as per the Carothers equation, or polymer precipitation

### Synthesis and Physicochemical Characterization of PCA-PEG Nanoparticles

PCA is a phenolic polymer derived from coumaric acid, a naturally occurring hydroxycinnamic acid found in fruits and vegetables such as tomatoes.^16,30^ In this study, PCA was co-assembled with DSPE-PEG during nanoprecipitation to form hybrid nanoparticles (NPs). DSPE-PEG, an amphiphilic polymer-lipid conjugate, facilitates NPs formation while providing steric stabilization and improves colloidal stability in aqueous media **(Figure 3a)**.^31,32^ To assess the effect of polymer structure on NPs formation, transmission electron microscopy (TEM) was performed on PCA-PEG NPs prepared from PCA polymers synthesized at different reaction times (0.5 h, 1 h, and 1.5 h), using fixed DSPE-PEG concentrations (5mg/mL). PCA0.5-PEG NPs exhibited irregular aggregation and poorly defined structures **(Figure 3c, S7)**. In contrast, PCA1-PEG NPs showed improved structural organization, with the emergence of a more distinct core **(Figure 3c, S8)**. Notably, PCA1.5-PEG NPs displayed a well-defined core-shell morphology with uniformly distributed particles **(Figure 3c, S9),** indicating more controlled NPs assembly. Dynamic light scattering (DLS) analysis of PCA1.5-PEG revealed a uniform hydrodynamic diameter (D_h_) of 109 nm and a polydispersity index (PDI) of 0.19 **(Figure 3d)**. To further optimize the NPs, the effect of DSPE-PEG stock was evaluated by preparing NPs 5, 10, and 20 mg/mL DSPE-PEG while keeping the polymer constant (PCA1.5h). The formulation containing 5 mg/mL DSPE-PEG (PCA1.5-PEG5 NPs) exhibited the most uniform size distribution, the lowest PDI, and a higher **ζ**-potential compared with PCA1.5-PEG10 and PCA1.5-PEG20 formulations (**Figure 3e, S6**). Therefore, 5 mg/mL DSPE-PEG was selected as the optimized PEG concentration for subsequent studies. For clarity, formulations evaluated during PEG optimization are denoted as PEG5, PEG10, and PEG20, corresponding to DSPE-PEG concentrations of 5, 10, and 20 mg/mL, respectively. After optimization, formulations prepared with 5 mg/mL DSPE-PEG are referred to as PCA-PEG or PCA-PEG5, unless otherwise specified. The reduced **ζ**-potential at higher DSPE-PEG concentrations could be attributed to increased surface shielding by the PEG corona.^33^ Although this near-neutral **ζ**-potential indicates that electrostatic repulsion is not the primary stabilizing mechanism, colloidal stabilization in these PEGylated formulations is expected to arise predominantly from PEG-mediated steric stabilization rather than charge-based repulsion.^34^ Finally, the colloidal stability of the optimized PCA1.5-PEG NPs was further assessed over 7 days under three different storage conditions (37 °C, 4 °C, and plasma). The NPs maintained consistent size distributions with minimal variations, indicating good colloidal stability under all tested conditions (**Figure 3f**).

**Figure 3.**
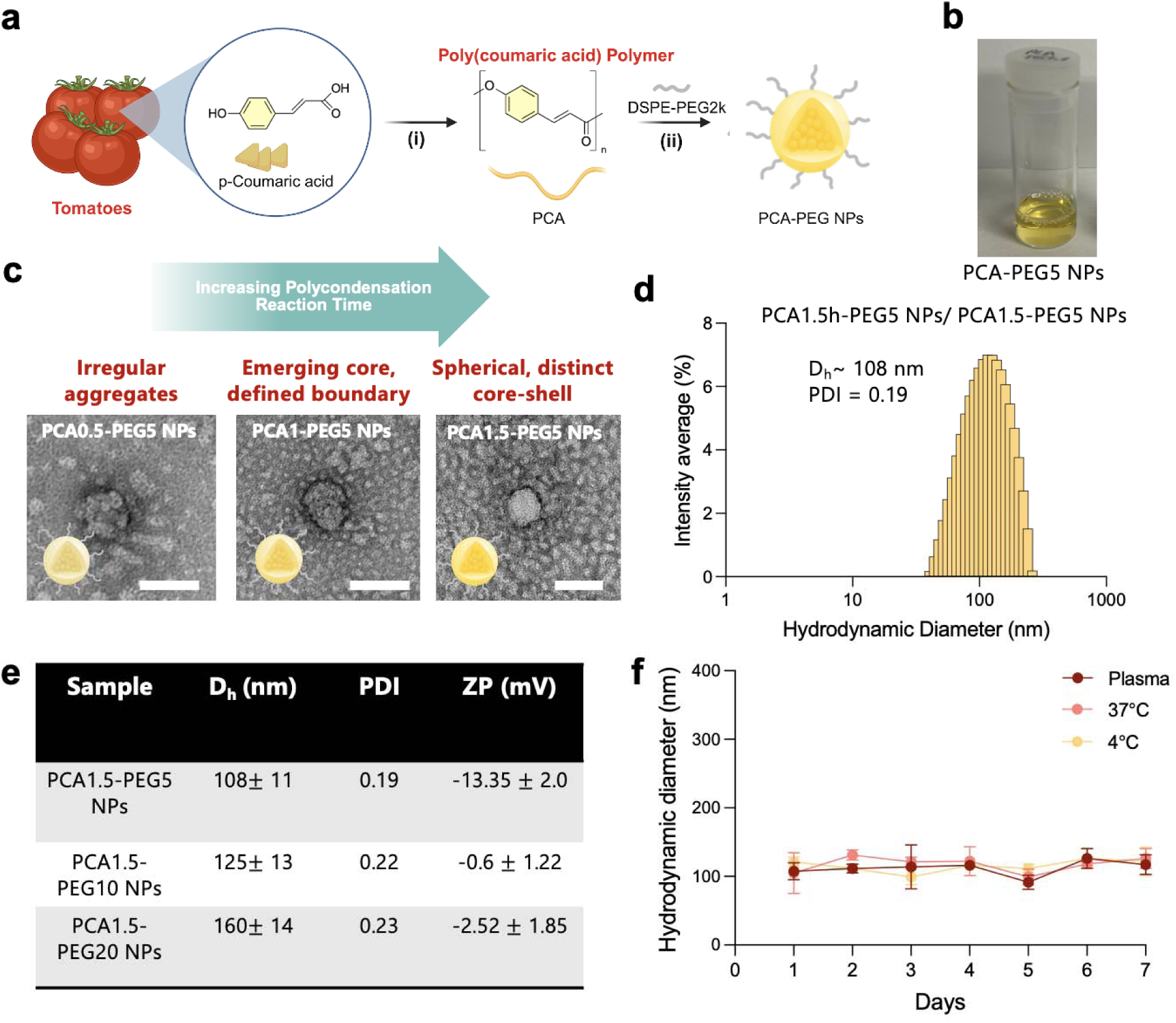
Synthesis, optimization, and physicochemical characterization of PCA-PEG NPs. (a) Schematic overview of PCA-PEG5 NPs synthesis using p-Coumaric acid polymer (b) Representative photograph of the formulated PCA1.5-PEG5 NPs (c) TEM images of PCA0.5-PEG5 NPs, PCA1-PEG5 NPs and PCA1.5-PEG5 NPs. Scale bar = 100 nm. (d) Hydrodynamic diameter intensity graph of PCA1.5-PEG NPs (e) Hydrodynamic diameter, PDI, and **ζ**-potential values of PCA-PEG5 NPs, PCA-PEG10 NPs and PCA-PEG20 NPs. (f) Colloidal stability of PCA1.5-PEG5 NPs over 7 days while incubated at 37°C, 4°C and 37°C with plasma. Data are presented as mean ± SD, n = 3.

Collectively, these findings suggest that reaction-time-dependent polymer growth influences nanoparticle assembly and stability, resulting in downstream differences in cargo encapsulation and biological performance. While the precise molecular mechanism remains to be fully elucidated, the observed trends establish a clear structure–property relationship between polycondensation conditions and nanoparticle function.

### Synthesis and Comprehensive Physicochemical and Optical Characterization of PCA-PEG-ICG NPs

Building on the optimized PCA1.5-PEG formulation, NPs were engineered to encapsulate indocyanine green (ICG), an FDA approved near-infrared (NIR) fluorescent dye, to impart imaging functionality.^23,35^ ICG was co-dissolved with PCA in DMSO and subsequently added dropwise into PBS under continuous probe sonication to form PCA-PEG-ICG NPs **(Figure 4a)**. Following purification *via* washing and filtration **(figure 4b)**, the physiochemical, optical properties and the encapsulation efficiency of the NPs were characterized. The DLS measurements revealed a hydrodynamic diameter of 127.2 nm and a PDI of 0.24, while the **ζ**-potential was approximately -11.3 mV **(Figure 4c, d)**. Compared with unloaded PCA1.5-PEG NPs, which exhibited a PDI of 0.19, ICG loading resulted in a modest increase in PDI to 0.24, indicating a slight broadening of the nanoparticle size distribution. This effect may arise from interactions between ICG and the PCA/DSPE-PEG matrix during nanoparticle self-assembly. Nevertheless, the resulting PDI remained below 0.3, consistent with a relatively uniform nanoparticle population. Although PCA1.5h-PEG-ICG NPs exhibited a moderately negative **ζ**-potential of approximately −11 mV, **ζ**-potential alone does not fully define colloidal stability, particularly for PEGylated nanoparticles. The DSPE-PEG component likely contributes PEG-mediated steric stabilization, consistent with the observed maintenance of hydrodynamic size and PDI under storage and plasma-containing conditions. Importantly, these properties were comparable to those of unloaded NPs, suggesting that ICG did not significantly alter the NPs characteristics. UV-Vis spectroscopy of PCA1.5-PEG-ICG NPs showed a characteristic peak ∼ 800 nm corresponding to encapsulated ICG **(Figure 4e)**, while fluorescence measurements (λ_ex_ = 785nm) demonstrated strong NIR-I emission with an emission maxima at 846nm **(Figure 4f)**. Fluorescence imaging *via* IVIS imaging further confirmed robust signal intensity compared to controls, with a concentration-dependent increase observed across serial dilutions **(Figure 4g, h)**. The encapsulation efficiency (EE) of ICG in PCA1.5-PEG-ICG NPs was 21.1%, exceeding that of PCA1-PEG-ICG NPs (17.6%) and PCA0.5-PEG-ICG NPs (13.6%). This increase in ICG encapsulation with longer PCA reaction time suggests that extended PCA polymer growth improves dye incorporation during nanoparticle self-assembly, likely by forming a more favorable hydrophobic/polymeric microenvironment for ICG retention **(Figure S10)**.

**Figure 4.**
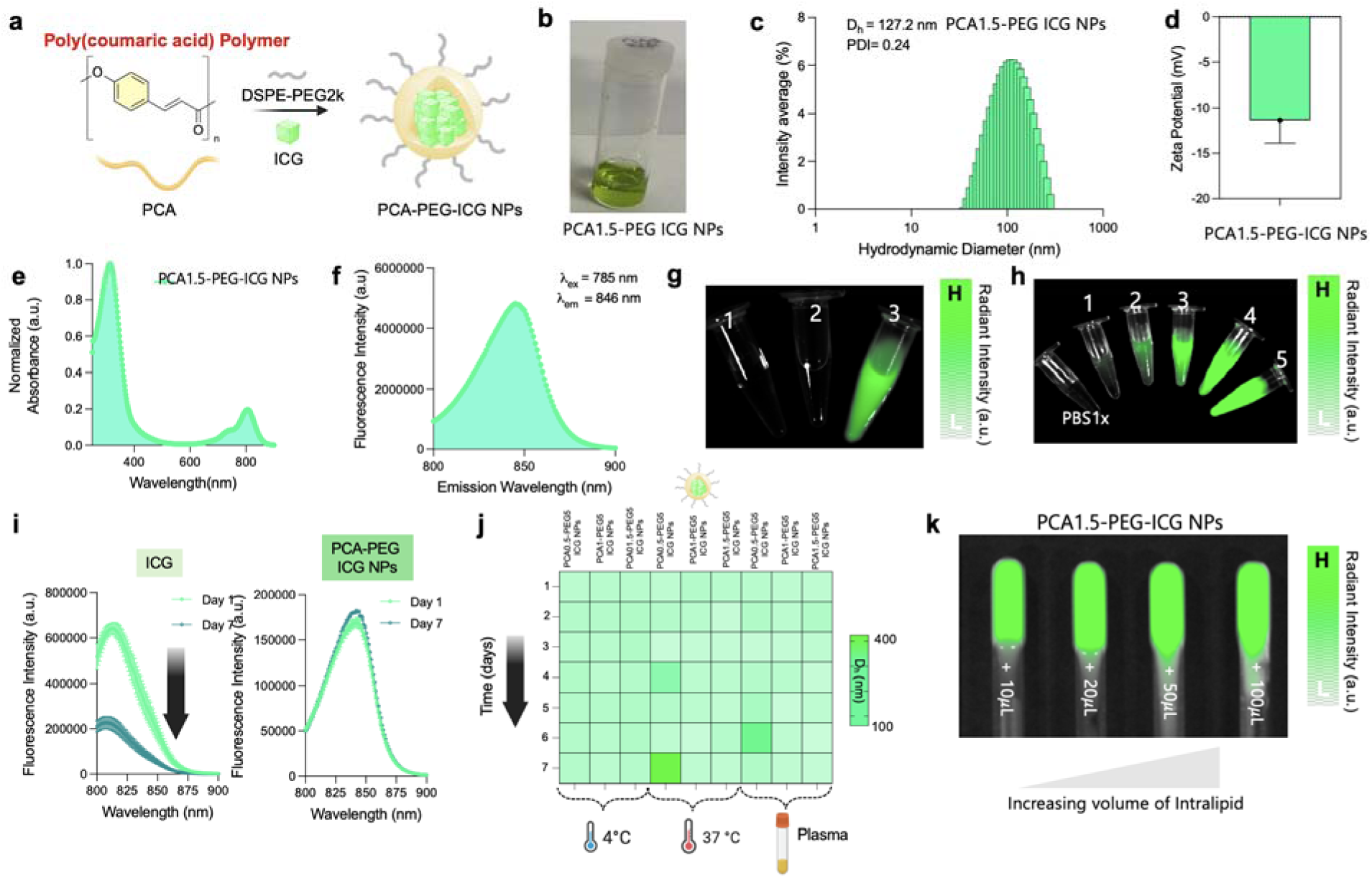
Preparation, physiochemical and optical characterization of PCA-PEG-ICG NPs. (a) Schematic overview of PCA-PEG-ICG NPs synthesis (b) Representative white light photograph of the formulated PCA1.5-PEG-ICG NPs (c) and (d) Hydrodynamic diameter intensity graph and Zeta potential (mV) of PCA-PEG-ICG NPs. (e) UV-VIS absorbance spectra of PCA-PEG-ICG NPs and (f) Fluorescence intensity of PCA-PEG-ICG NPs. (g) NIR-I fluorescence imaging of (1) PBS-1X (2) PCA1.5-PEG NPs (3) PCA1.5-PEG-ICG NPs. (h) NIR-I fluorescence imaging of PBS1X and PCA-PEG-ICG NPs, where 5: 7.3 **μ**M , 4: 3.5 **μ**M, 3: 1.9 **μ**M, 2: 0.69 **μ**M and 1: 0.12 **μ**m of ICG. (i) Fluorescence stability of PCA-PEG-ICG NPs and free ICG evaluated on day 1 and day 7. (j) Colloidal stability of PCA0.5-PEG-ICG NPs, PCA1-PEG-ICG NPs and PCA1.5-PEG-ICG NPs over 7 days while incubated at 37°C, 4°C and 37°C with plasma. (k) NIR-I fluorescence imaging of PCA1.5-PEG-ICG NP in an NMR tube with increasing volume of intralipid.

Given the known limitations of free ICG, including poor photostability and aggregation in aqueous environments,^36^ we next evaluated fluorescence stability over time. PCA-PEG-ICG NPs maintained stable fluorescence signal intensity over a period of 7 days, whereas free ICG exhibited substantial signal decay, highlighting the protective role of nanoparticle encapsulation **(Figure 4i)**. Quantitative comparison of Day 1 and Day 7 fluorescence intensity confirmed that free ICG underwent a significant decrease in fluorescence over 7 days, while PCA1.5-PEG-ICG NPs showed no significant reduction over the same period **(Figure S11)**.

To distinguish the contribution of the PCA-containing matrix from DSPE-PEG-mediated encapsulation alone, we also evaluated PEG-ICG NPs lacking PCA. PEG-ICG NPs retained nanoparticle-associated fluorescence over 7 days, indicating that DSPE-PEG encapsulation contributes to ICG stabilization **(Figure S12)**. However, comparison of free ICG, PEG-ICG NPs, and PCA1.5-PEG-ICG NPs on Day 7 showed that PCA1.5-PEG-ICG NPs provided the most stable fluorescence signal among the tested formulations **(Figure S13)**. These results suggest that while nanoencapsulation improves ICG stability, the PCA-containing hybrid matrix provides additional stabilization beyond PEG-ICG encapsulation alone.

Colloidal stability was further assessed by comparing PCA 1.5-PEG-ICG NPs with lower polymer formulations (PCA1-PEG-ICG NPs and PCA0.5-PEG-ICG-NPs) under physiological (37°C), 37°C with plasma, and refrigerated (4°C) for 7 days. PCA1.5-PEG-ICG showed minimal fluctuations in hydrodynamic size across all conditions, indicating high colloidal stability across all conditions, indicating improved colloidal stability relative to other formulations **(Figure 4j)**. Furthermore, incubation in plasma resulted in negligible changes in particle size, indicating favorable interactions with serum proteins and resistance to protein-induced aggregation. Such behavior could be attributed to the increased polymer length, leading to a higher encapsulation efficiency of PCA1.5-PEG-ICG NPs in comparison to its counterparts **(Figure S10)**, reducing ICG aggregation and providing protection against photodegradation.

To further evaluate cargo retention, ICG release from PCA1.5-PEG-ICG NPs was assessed under physiological (pH 7.4) and acidic (pH 5.0) conditions using a dialysis-based release assay. Dialysate fractions were collected over 24 h and analyzed by UV–Vis absorbance and fluorescence spectroscopy, while the nanoparticle retentate was analyzed at 0 and 24 h. Under both pH conditions, the dialysate fractions showed minimal characteristic ICG absorbance and only weak fluorescence signal relative to the retentate, indicating limited detectable ICG release over 24 h (Figures S14 and S15). In contrast, the retentate maintained the characteristic optical features of encapsulated ICG, supporting strong dye retention within PCA1.5-PEG-ICG NPs under both physiological and acidic conditions.

Because the dialysate fractions showed minimal ICG-associated absorbance and fluorescence under both pH conditions, the difference in retentate fluorescence is not consistent with substantial dye release. Rather, the lower retentate fluorescence observed at pH 5.0 indicates reduced fluorescence efficiency of retained nanoparticle-associated ICG under acidic conditions. In contrast, the stronger retentate fluorescence at pH 7.4 suggests better preservation of the optical properties of encapsulated ICG under physiological conditions. Importantly, because the dialysate showed only minimal ICG-associated signal under both conditions, the lower retentate fluorescence at pH 5.0 is more consistent with reduced fluorescence efficiency of retained nanoparticle-associated ICG than with substantial dye release. The stronger retentate fluorescence at pH 7.4 further supports the optical stability of PCA1.5-PEG-ICG NPs under physiological conditions.

To better approximate the optical complexity of tumor environments, fluorescence performance was evaluated in the presence of intralipid, a scattering medium that mimics soft tissue. Notably, increasing intralipid concentration did not significantly attenuate fluorescence intensity, indicating strong signal retention under highly scattering conditions **(Figure 4k, S16)**. Furthermore, evaluation within TMPs composed of intralipid and hemoglobin **(Figures S25, S26)** demonstrated detectable fluorescence even at low concentrations, supporting the suitability of PCA-PEG-ICG nanoparticles for imaging in complex tissue-like environments.

### Assessment of *In Vitro* Cytotoxicity, Cellular Internalization and Biological Activity of PCA-PEG and PCA-PEG-ICG NPs

Coumaric acid possesses intrinsic cytotoxic and oxidative stress-modulating properties.^20,30,37^ We therefore hypothesized that polymerizing coumaric acid into PCA would generate an active phenolic scaffold that could be formulated into stable PEGylated NPs while retaining biologically relevant redox activity. Because reaction time altered PCA polymer growth and nanoparticle assembly, we further evaluated whether these structural differences translated into formulation-dependent cytotoxicity. To test this, we compared the cytotoxic profiles of PCA0.5-PEG, PCA1-PEG, and PCA1.5-PEG NPs in OVCAR8 cells. As shown in **Figure 5a, b**, PCA1.5-PEG NPs produced a greater concentration-dependent reduction in OVCAR8 cell viability compared with the lower reaction-time formulations, PCA0.5-PEG and PCA1-PEG. A similar concentration-dependent decrease in cell viability was observed in OVCAR3 cells following treatment with PCA-PEG NPs, supporting activity across an additional ovarian cancer cell line **(Figure S17a–c)**. In contrast, PEG5 NPs lacking PCA did not induce a concentration-dependent decrease in cell viability, supporting the contribution of the PCA component to the observed biological response **(Figure S18)**. We also evaluated normal-cell toxicity using HUVECs. PCA1.5-PEG NPs did not produce a comparable decrease in HUVEC viability up to 50 **μ**g/mL, suggesting reduced toxicity toward non-cancerous endothelial cells under the tested conditions **(Figure S19)**.

**Figure 5.**
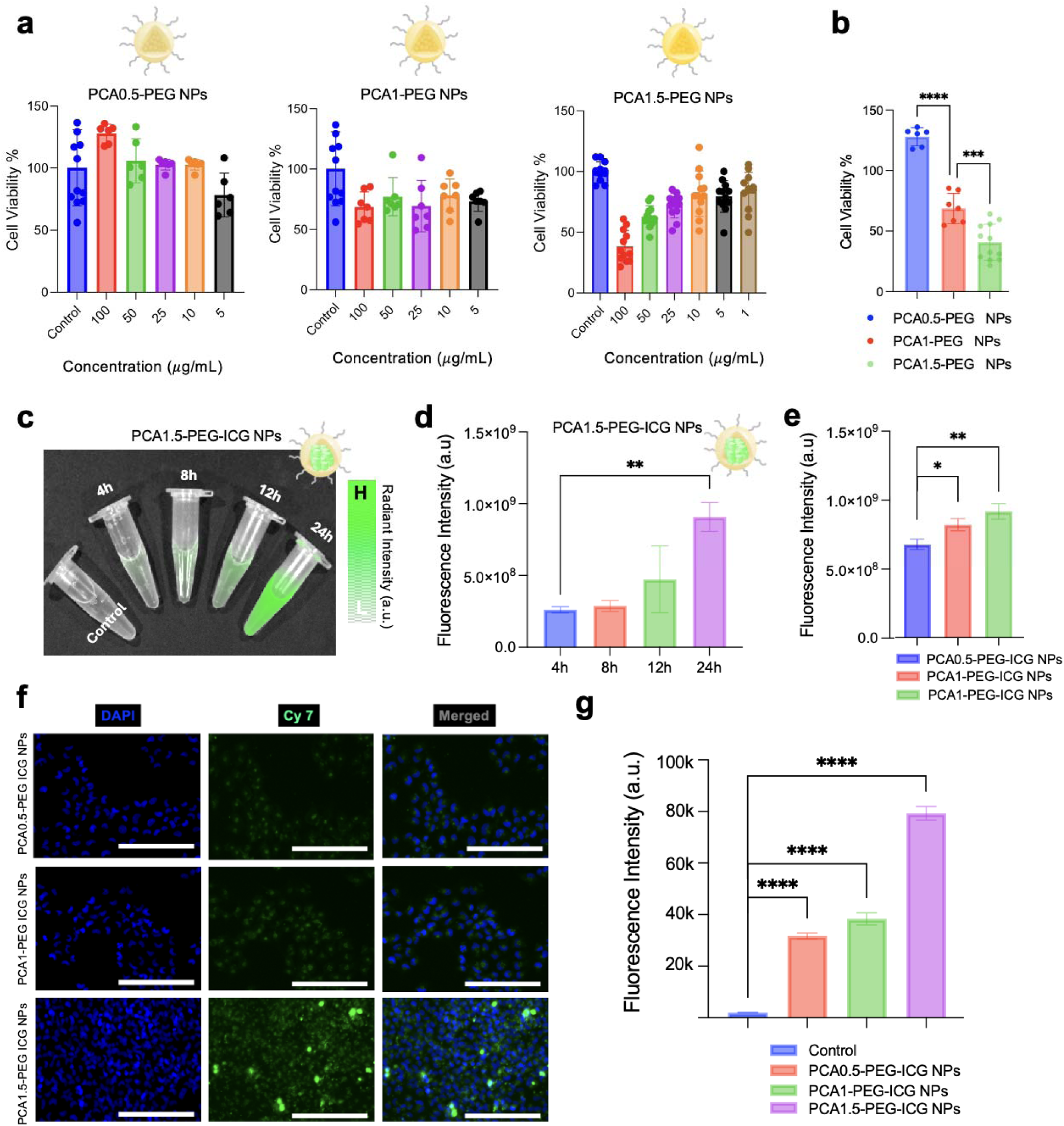
*In-vitro* evaluation of cytotoxicity, cellular uptake, and biological activity of PCA-PEG NPs. (a) Cell viability of OVCAR8 following treatment with PCA0.5-PEG NPs, PCA1-PEG NPs and PCA1.5-PEG NPs. (b) Comparative analysis of cell viability in OVCAR8 following treatment with 100 μg/ml PCA0.5-PEG NPs, PCA1-PEG NPs and PCA1.5-PEG NPs. (c) NIR-I fluorescence imaging of OVCAR8 following treatment with PCA1.5-PEG-ICG NPs at 4h, 8h, 12 and 24h. (d) Quantification of fluorescence intensity of OVCAR8 following treatment with PCA1.5-PEG-ICG NPs at 4h, 8h, 12 and 24h showing a time dependent internalization. (e) Quantification of fluorescence intensity of OVCAR8 following treatment with PCA0.5-PEG-ICG NPs, PCA1-PEG-ICG NPs and PCA1.5-PEG-ICG NPs at 24h. (f) Representative fluorescence microscopy images of OVCAR8 cells following treatment with PCA0.5-PEG NPs, PCA1-PEG NPs and PCA1.5-PEG NPs showing cellular internalization at 24 hours. (g) Quantification of fluorescence intensity of OVCAR8 following treatment with PCA0.5-PEG-ICG NPs, PCA1-PEG-ICG NPs and PCA1.5-PEG-ICG NPs at 24h.

As nanoparticle–cell association can influence biological response, we evaluated PCA-PEG-ICG NP uptake in OVCAR8 and OVCAR3 cells using IVIS imaging and complementary fluorescence microscopy **(Figure 5c–g and Figure S20 a,b)**. Based on their stronger cytotoxic response, PCA1.5-PEG-ICG NPs were first selected for time-dependent uptake profiling. Fluorescence quantification showed a time-dependent increase in cell-associated PCA1.5-PEG-ICG NP signal, with the highest fluorescence observed at 24 h **(Figure 5c,d and Figure S20a,b).** Because nanoparticle–cell interactions are influenced by physicochemical properties such as morphology, surface charge, and colloidal stability,^33,38–40^ we next compared PCA0.5-PEG-ICG, PCA1-PEG-ICG, and PCA1.5-PEG-ICG NPs at the 24 h time point. PCA1.5-PEG-ICG NPs exhibited the highest cell-associated fluorescence among the tested formulations by IVIS-based quantification **(Figure 5e)**.Complementary fluorescence microscopy further supported this trend, showing stronger Cy7-channel fluorescence in cells treated with PCA1.5-PEG-ICG NPs compared with lower reaction-time formulations **(Figure 5f,g)**. These results suggest that reaction-time-dependent differences in PCA growth profile and nanoparticle assembly influence cell-associated nanoparticle fluorescence. However, the present experiments do not define a specific internalization pathway or subcellular trafficking route.

To further elucidate the biological mechanism underlying PCA1.5-PEG NP-induced cytotoxicity, intracellular ROS generation was assessed using the H2DCFDA fluorescent probe (**Figure 6a**). Representative fluorescence microscopy images showed increased DCF fluorescence in PCA1.5-PEG NP-treated OVCAR8 and OVCAR3 cells relative to untreated controls, with signal intensity increasing as a function of nanoparticle concentration and treatment duration **(Figure 6b–d and Figures S21)**. Quantitative analysis confirmed a significant increase in ROS-associated fluorescence following exposure to PCA1.5-PEG NPs at 2, 4, and 8 h, with the highest signal observed after 8 h treatment at 50 **μ**g/mL **(Figure 6e–g and Figures S21)**. These results indicate that PCA1.5-PEG NPs induce time- and concentration-dependent oxidative stress in ovarian cancer cells.

**Figure 6.**
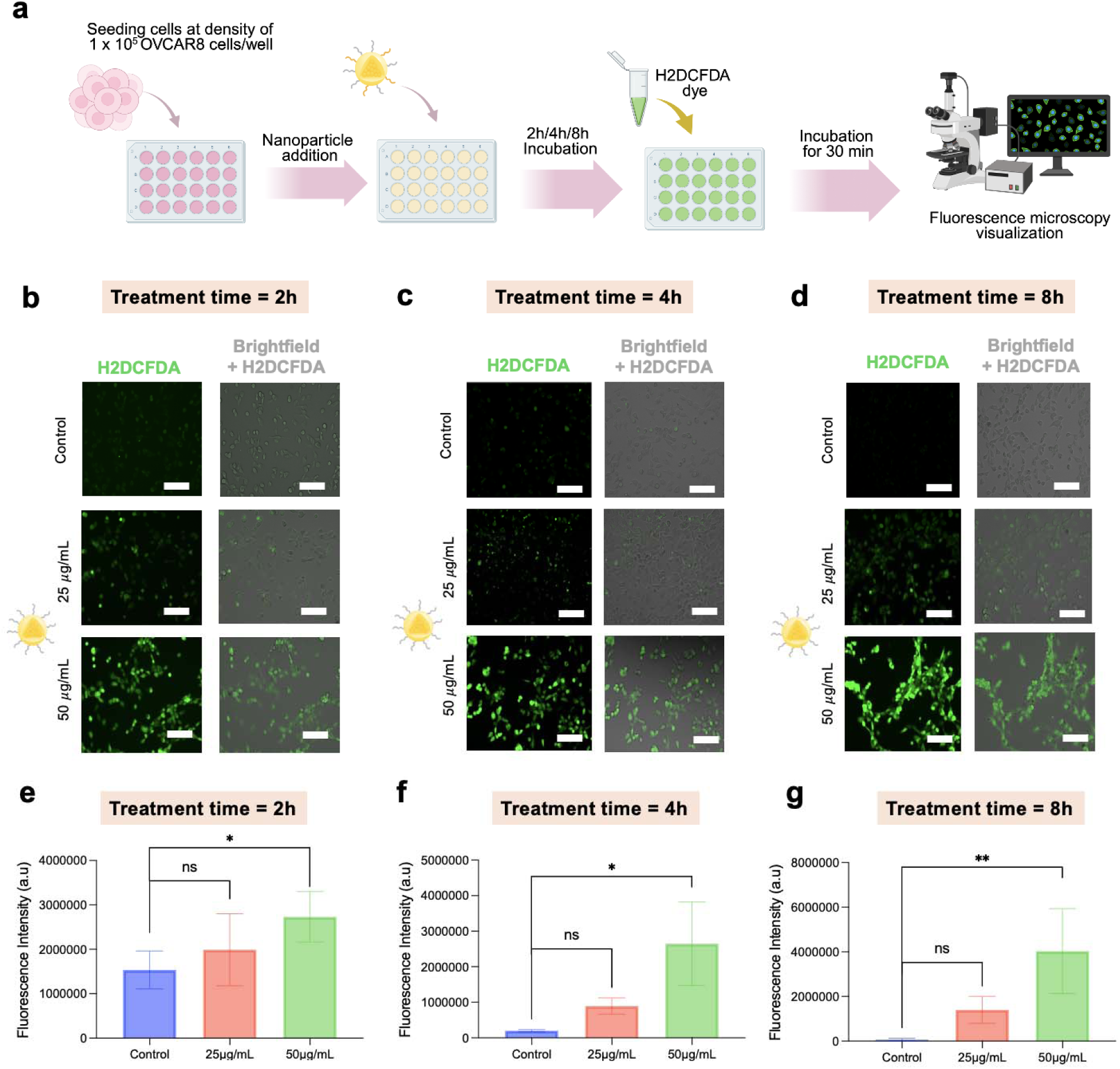
*In vitro* assessment of intracellular reactive oxygen species (ROS) generation following PCA1.5-PEGnanoparticle treatment in OVCAR8 cells. (a) Schematic illustration of the experimental workflow used for ROS detection. OVCAR8 cells were seeded in 24-well plates at a density of 1 × 10O cells/well and allowed to attach for 24 h prior to treatment with PCA1.5-PEG nanoparticles. Cells were exposed to nanoparticles at the indicated concentrations for 2, 4, or 8 h, followed by incubation with 20 μM H2DCFDA for 30 min at 37°C. Fluorescence images were then acquired using an EVOS M5000 fluorescence microscope. Representative fluorescence microscopy images showing intracellular ROS levels (green fluorescence, H2DCFDA) after treatment with PCA1.5-PEG nanoparticles for (b) 2 h, (c) 4 h, and (d) 8 h. Corresponding merged brightfield and fluorescence images are shown alongside each treatment group. Control cells received no nanoparticle treatment. Quantification of fluorescence intensity following nanoparticle exposure for (e) 2 h, (f) 4 h, and (g) 8 h demonstrated a time- and concentration-dependent increase in ROS production, with higher fluorescence intensity observed in nanoparticle-treated groups compared with control cells. Data are presented as mean ± SD (n = 4). Statistical significance is indicated as ns, not significant; *p < 0.05; **p < 0.01. Scale bars represent 150 **μ**m.

These findings indicate that PCA1.5-PEG NPs retain oxidative stress-modulating bioactivity following nanoparticle formulation. Elevated ROS levels can disrupt cellular redox homeostasis and contribute to cancer cell death through multiple downstream pathways.^41^ Therefore, the observed increase in H2DCFDA fluorescence is consistent with the cytotoxic effects of PCA1.5-PEG NPs observed in the MTT assay. However, the present data do not define a specific apoptotic, mitochondrial, or other downstream cell-death pathway. Accordingly, we interpret these results as evidence of ROS-associated cytotoxicity rather than definitive ROS-mediated apoptosis.

Collectively, these results establish PCA1.5-PEG NPs as the most effective formulation among the tested candidates, combining superior cellular uptake, potent cytotoxicity, and robust ROS-mediated biological activity. The convergence of these properties highlights the importance of polymer chain length and nanoparticle architecture in governing intracellular delivery performance and therapeutic efficacy. Moreover, the strong uptake profile and preserved biological function of PCA1.5-PEG-ICG NPs support their continued development as a theranostic nanoplatform for ovarian cancer treatment.

### Photobleaching and Photostability Evaluation of PCA-PEG-ICG Nanoparticles

Fluorescence based imaging is frequently limited by photobleaching, particularly for clinically used fluorophores such as ICG, which undergo rapid signal degradation under continuous illumination. This primarily occurs due to the photoinduced interactions with molecular oxygen and reactive species that result in irreversible structural degradation.^42–44^

Photostability of our PCA-PEG-ICG NPs was evaluating tumor-mimicking phantoms (TMPs), previously developed by us, ^23–25,45^ which more closely recapitulates the optical and absorption characteristics of tumors compared to conventional *in vitro* measurements. These phantoms incorporate selected tissue-relevant optical components, including absorbers and scatterers, to enable reproducible evaluation of fluorescence signal retention and attenuation under defined tumor-mimicking conditions. Importantly, these models are not intended to reproduce vascular perfusion, immune interactions, nanoparticle biodistribution, or the full heterogeneity of *in vivo* tumors, but rather to provide a preclinical optical testbed prior to more complex *in vivo* studies.

Upon continuous white light excitation (1000 lumens) (**Figure S27**), PCA-PEG-ICG NPs TMPs retained strong fluorescence over a 4h period, whereas free ICG at similar concentrations exhibited rapid signal decay under identical conditions (Figure 7a-c). Control TMPs without any contrast agents showed negligible signal. To determine whether this enhanced photostability was attributable solely to DSPE-PEG-mediated nanoencapsulation, we evaluated DSPE-PEG-ICG TMPs lacking PCA under the same illumination and imaging conditions **(Figure S27)**, DSPE-PEG-ICG TMPs showed partial protection relative to free ICG TMPs **(Figure S28)**; however, fluorescence decay was more pronounced than in PCA1.5-PEG-ICG TMPs, particularly after prolonged illumination. These findings indicate that DSPE-PEG nanoencapsulation contributes to ICG stabilization, while the PCA-containing hybrid matrix provides additional protection, likely by creating a more protective hydrophobic/phenolic microenvironment that reduces dye aggregation and photodegradation. Quantitative radiant efficiency analysis confirmed improved fluorescence retention for PCA1.5-PEG-ICG TMPs relative to both free ICG and DSPE-PEG-ICG controls. The enhanced photostability can be attributed to multiple synergistic effects of the PCA matrix. First, the polymeric network acts as a physical barrier that limits oxygen diffusion and reduces direct interaction of excited ICG with external quenchers.^46–49^ Second, the phenolic functionality of PCA provides intrinsic antioxidant activity, enabling scavenging of reactive oxygen species generated during photoexcitation and thereby suppressing oxidative degradation pathways.^18, 20, 26^ Additionally, encapsulation within the hydrophobic polymer matrix reduces dye aggregation and restricts conformational flexibility, helping to maintain ICG in a more stable and dispersed state.^46^

**Figure 7.**
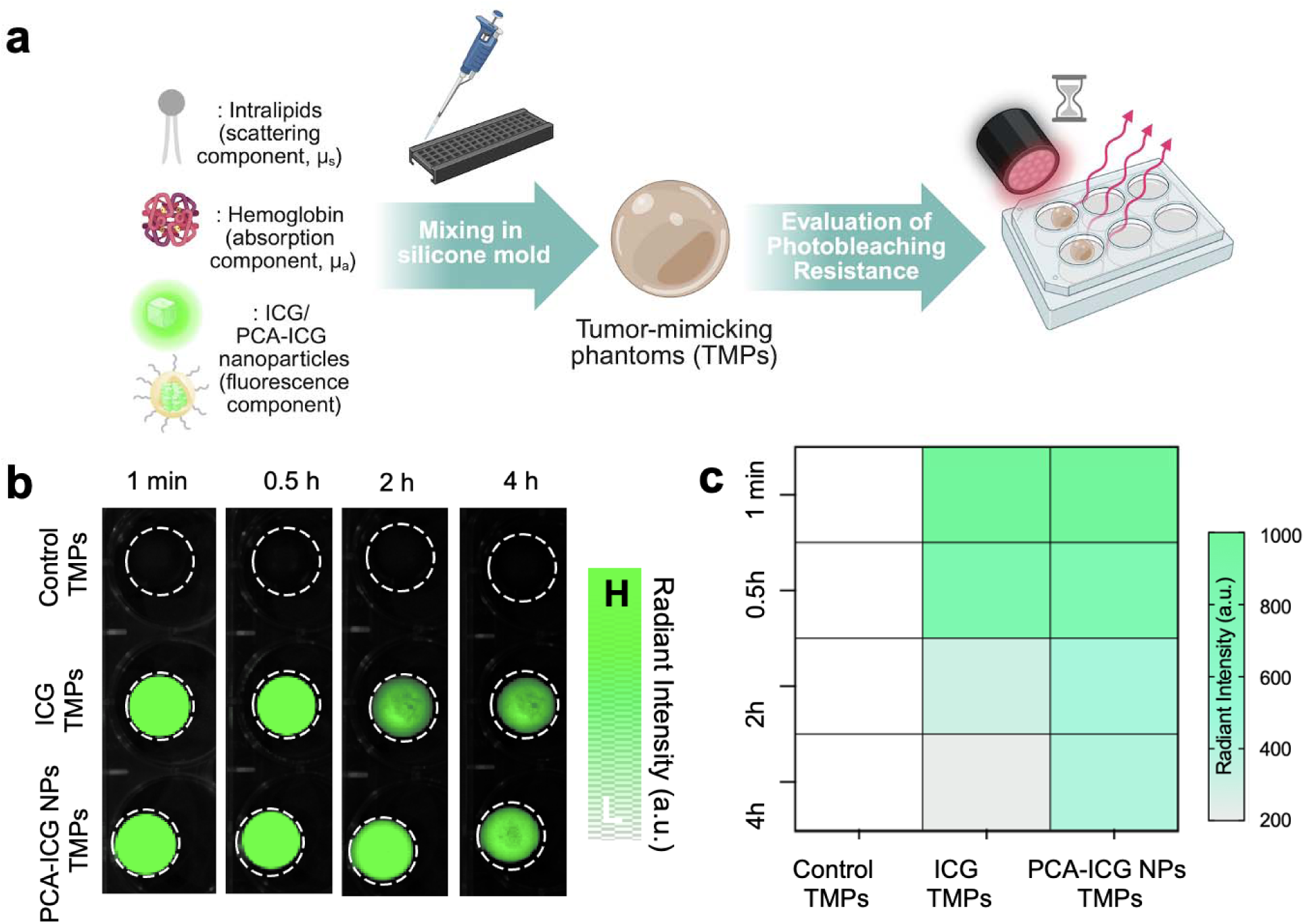
Photobleaching assessment of PCA-PEG-ICG NP TMPs compared to clinical standard, ICG. (a) Schematic diagram of preparation of PCA-PEG-ICG NPs and ICG TMPs and their photostability evaluation. (b) NIR-I fluorescence image of Control TMP, PCA-ICG NP TMP and ICG TMP after being subject to 1000 lumen white light source for different time durations (1min, 0.5h, 2h, 4h). (c) Corresponding radiant fluorescence intensity measurements for (b). Experiments were done in triplicates.

Tissue penetration profiling under clinically relevant settings was further performed to evaluate the optical performance of PCA-PEG-ICG NPs. Layered *ex vivo* porcine tissues (muscle, fat, and skin) were used to simulate relevant optical barriers and varying penetration depth (0, 2, 4 mm) **(Figure S29)**. Compared to ICG TMPs at equivalent concentrations, PCA-PEG-ICG NPs TMPs consistently exhibited higher fluorescence intensity across all tissue types and increasing depths **(Figure S30-S32)**. The enhanced signal propagation through tissue is attributed to the improved stability and preserved fluorescence of ICG within the PCA matrix, which mitigates signal loss under attenuating conditions. These results further highlight the ability of the PCA-based nanoparticle system to maintain fluorescence output in optically challenging environments, supporting its potential for improved intraoperative imaging, particularly in deeper or highly scattering tissues.

### Performing Surgical Simulation on 3D-printed Intraperitoneal Tumor Phantoms

Building on our previously reported 3D tumor-mimicking phantoms (TMPs) platform, ^23–25,45^ which enables precise control over tissue optical properties and tumor composition through a customizable bioprinted bioink, we sought to extend this system to more clinically relevant cancer scenarios.

Intraperitoneal (IP) tumors represent a particularly challenging and clinically relevant model for evaluating image-guided surgical approaches due to their diffuse anatomical distribution, the frequent requirement for extensive cytoreductive surgery (CRS) to remove multiple scattered nodules, and a heterogeneous microenvironment that limits the effectiveness of conventional therapies.^50–53^

While our prior TMPs constructs effectively recapitulated the optical and structural features of individual tumors, they did not capture the multifocal and spatially heterogeneous nature of IP metastases,^54–56^ where numerous tumor nodules are distributed throughout the abdominal cavity. To address this limitation, we developed a 3D bioprinted IP tumor model incorporating multiple TMPs within a tissue mimicking-environment to replicate the nodule dissemination observed within an IP tumor-bearing mice (Figure 8a). To further approximate physiological imaging conditions, we fabricated subcutaneous skin-mimicking layers with tunable melanin concentrations **(Figure S33)** to replicate the optical attenuation and scattering imposed by overlaying tissues, which subsequently influences fluorescence signal propagation during *in vivo* imaging of IP tumors.

**Figure 8.**
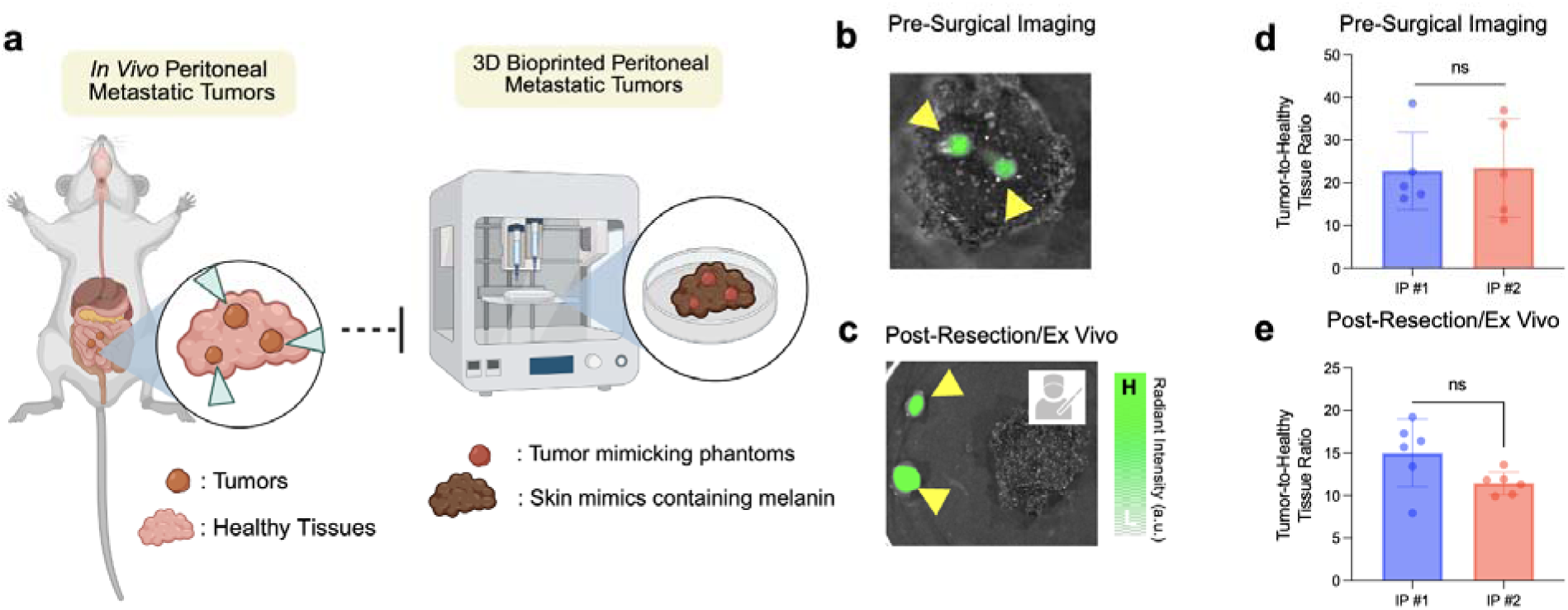
3D-bioprinted intraperitoneal tumor model for simulated ex vivo image-guided surgery. (a) Schematic showing the design of a 3D-bioprinted intraperitoneal tumor-mimicking model inspired by multifocal peritoneal metastatic disease. Tumor-mimicking phantoms containing PCA1.5-PEG-ICG NPs were printed onto melanin-containing skin-mimicking constructs. Representative NIR-I fluorescence images of (b) pre-surgical imaging and (c) post-resection/ex vivo imaging showing fluorescent tumor-mimicking regions. Quantification of tumor-to-healthy tissue fluorescence ratio for (d) pre-surgical imaging and (e) post-resection/ex vivo imaging. IP #1 and IP #2 represent independently printed constructs generated using the same phantom composition and printing parameters. Each datapoint represents an individual tumor-to-healthy tissue region-of-interest measurement from the corresponding printed construct. Data are presented as mean ± SD, n = 6, ROI measurements per construct. Statistical comparisons between IP #1 and IP #2 were performed using an unpaired two-tailed Welch’s t-test; ns, not significant.

TMPs were generated using our established bioink formulation containing optical scatterers (Intralipid), absorbers (hemoglobin), and a gelatin-based biomimetic matrix, along with PCA-PEG-ICG NPs as the functional imaging agent. This system enables precise control over tumor geometry, composition, and spatial distribution within a precisely controlled matrix. The printed constructs were integrated into a controlled, tumor-mimicking IP phantom environment **(Figure S34).** Rather than representing a fixed optical model, the TMP platform was designed as a modular system in which key optical and structural parameters can be independently varied. Hemoglobin was used as an absorber to modulate absorption-associated fluorescence attenuation, Intralipid was used as a scattering component to alter fluorescence propagation, and melanin was incorporated into skin-mimicking layers to model pigmentation-dependent attenuation. Supporting experiments demonstrate that increasing hemoglobin, Intralipid, and melanin content alters the optical behavior and fluorescence readout of PCA1.5-PEG-ICG NPs **(Figure. S22-S24).** In addition, stacked tissue layers produced depth-dependent signal attenuation **(Figure S30-S32)**, while melanin-containing bioprinted skin mimics enabled construction of skin-like layers with variable **pigmentation (Figure S33)**. These tunable components were then incorporated into 3D-bioprinted skin/IP tumor constructs containing multiple spatially distributed TMPs **(Figure S34).**

Fluorescence-guided surgical resection was then performed to simulate image-guided surgery, enabling real-time visualization of tumor margin and assessment of nanoprobe distribution and contrast during resection (Figure 8a-d). This platform allowed systematic evaluation of NPs performance under dynamic settings, including tissue-like scattering, repeated light exposure, and partial resections that mimic intraoperative procedures. Importantly, residual tumor-to-background signal post-resection was quantified to assess the ability of NPs to identify incompletely removed tumor foci **(Figure 8c, e)**. Consistent with our earlier TMP results showing rapid photobleaching of free ICG TMPs under comparable illumination conditions (Figure 7b), PCA1.5-PEG-ICG NPs maintained robust fluorescence contrast within the 3D-bioprinted IP model. Although this platform does not replace in vivo biodistribution, pharmacokinetic, tumor accumulation, or head-to-head comparative studies, it provides a reproducible preclinical testbed for evaluating nanoparticle fluorescence contrast, tumor delineation, and simulated image-guided resection under defined tissue-mimicking optical conditions. IP #1 and IP #2 represent independently printed intraperitoneal tumor constructs generated using the same tumor-mimicking phantom composition, skin-mimicking composition, and printing parameters (Fig. 8b**–e**). These constructs were used to assess whether fluorescence contrast measurements were consistent across separate prints of the same model design.

Overall, this 3D bioprinted platform provides a reliable and reproducible method for studying NPs-assisted tumor delineation, intraoperative decision-making, and resection accuracy in a preclinically relevant imaging context, effectively bridging the gap between simplified in vitro assessments and complex in vivo studies.

### Hemocompatibility Assessments of PCA-PEG NPs

Assessing how nanoprobes interact with red blood cells is a critical early step in determining their biocompatibility.^24,45^, In this study, we measured hemolysis, defined as the rupture of erythrocytes, after incubating porcine red blood cells with PCA-PEG nanoparticles at varying concentrations (1, 5, 10, 50, and 100 **μ**g/mL). Standard controls were included for comparison: phosphate-buffered saline (PBS1x) served as the negative control, while deionized water acted as the positive control. All samples treated with PCA-PEG nanoparticles showed intact red blood cell pellets at the bottom of the tubes, indicating that cell integrity was maintained (Figure 9a**, c**). In contrast, samples exposed to deionized water exhibited complete hemolysis, as shown by a uniformly red supernatant and the absence of a pellet. The PBS1x control produced a compact pellet with a clear supernatant, consistent with minimal hemolysis (Figure 9b). Across all tested concentrations, PCA-PEG nanoparticles induced hemolysis levels below the 10% threshold considered acceptable for blood compatibility, with the lowest value observed at 10 **μ**g/mL (3.12 ± 1.08%) (Figure 9d). These findings demonstrate that the nanoprobes are hemocompatible within the tested concentration range and are unlikely to cause red blood cell damage under physiological conditions.

**Figure 9.**
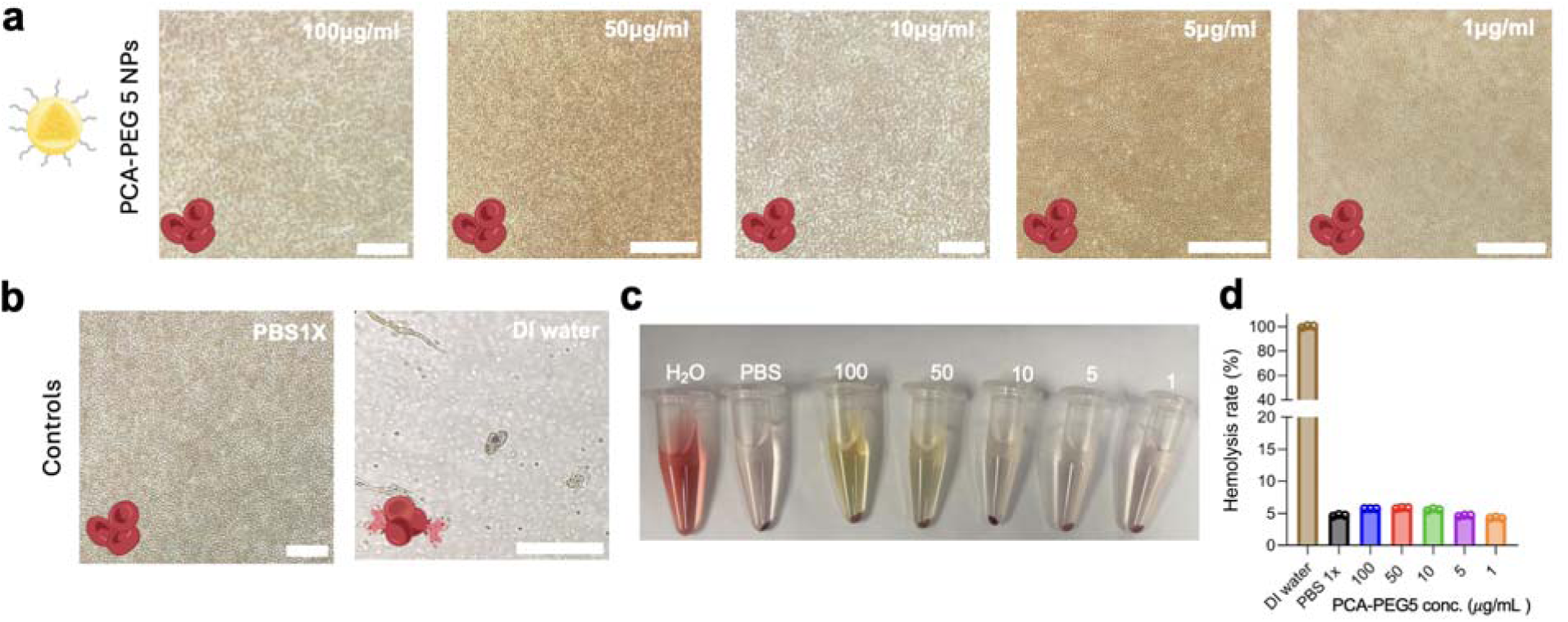
Hemocompatibility studies of PCA-PEG NPs with porcine blood. Inverted microscopic images at 200× magnification of (a) PCA-PEG NPs (100, 50, 10, 5, and 1 μg/mL), PBS1x (negative control), and DI water (positive control) after 1h of incubation at 37 °C with porcine blood. (c) White light image of RBC pellets from each incubated sample: (1) 100 μg/mL, (2) 50 μg/mL, (3) 10 μg/mL, (4) 5 μg/mL, (5) 1 μg/mL, (6) PBS1x, and (7) DI water. (d) Relative hemolysis percentages of all PCA-PEG treatment and control groups.

Overall, these preliminary results suggest minimal toxicity in healthy porcine blood at the evaluated doses, supporting their potential use for *in vivo* applications. Nonetheless, more comprehensive studies are needed to assess both short- and long-term toxicity across a wider range of doses in living systems, which falls beyond the scope of this work.

## CONCLUSIONS

In summary, we report the development of self-theranostic hybrid nanoparticles in which the macromolecular architecture of the PCA core serves as a key determinant of both optical and colloidal performances, cellular endocytosis behavior, and therapeutic efficacy. By tuning the polymer structure through variation in polycondensation reaction time, we demonstrate that extended reaction durations promote the formation of more robust polymer backbones, enabling improved core-shell assembly. This optimized architecture enhances the hydrophobic sequestration and photostability of encapsulated ICG, a representative NIR-I fluorophore, effectively mitigating the rapid quenching and degradation associated with free dye systems.

Beyond its structural role, the PCA matrix functions as a redox-active, self-therapeutic scaffold. The nanoparticles exhibit preferential internalization in ovarian cancer cells, where the phenolic backbone modulates intracellular oxidative stress, ultimately inducing apoptosis. The dual imaging-therapeutic functionality was further validated using 3D bioprinted intraperitoneal tumor models, where the nanoprobes enabled high-contrast and photostable fluorescence for surgical guidance under optically complex conditions. Overall, this work establishes reaction-time-dependent PCA growth profiles as a critical design parameter associated with nanoparticle assembly, imaging performance, and biological activity.

### AUTHOR INFORMATION

## Corresponding Author

*

## Author Contributions

+ M. M., A. H., and K. H. contributed equally. I. S. conceived the idea and designed all the experiments. M. M., A. H., and K. H. were involved in data collection and curation, formal analysis, investigation, methodology, validation, and visualization of all the experiments under the supervision of I.S. 3D bioprinting of TMPs were conducted by I. D. L. under the supervision of I.S. M.H.R. performed transmission electron microscopy images. GPC was conducted by R.P. under the supervision of J.T. The first draft of the manuscript was written by M.M., A.H., K.H. and I.S. with edits from all the authors. All authors have approved the final version of the manuscript.

### ASSOCIATED CONTENT

## Supporting Information

Experimental data includes PCA polycondensation synthesis scheme (Figure S1); ^1^H-NMR characterization of CA and PCA polymers at different reaction times (Figures S2-S5); synthesis, optimization, hydrodynamic diameter, PDI, **ζ**-potential, and physicochemical characterization of PCA1.5-PEG NPs (Figure S6); negative-stained TEM images of PCA0.5-PEG, PCA1-PEG, and PCA1.5-PEG NPs (Figures S7-S9); ICG encapsulation efficiency in PCA-PEG-ICG NPs (Figure S10); fluorescence stability of free ICG, PEG-ICG NPs, and PCA1.5-PEG-ICG NPs over 7 days (Figures S11-S13); in vitro ICG release assessment of PCA1.5-PEG-ICG NPs under acidic and physiological pH conditions (Figures S14-S15); effect of increasing intralipid concentration on the optical performance of PCA-PEG-ICG NPs (Figure S16); in vitro cytotoxicity of PCA-PEG NPs, PEG5 NPs, and PCA1.5-PEG NPs in OVCAR3, OVCAR8, and HUVECs (Figures S17-S19); time-dependent cellular uptake of PCA1.5-PEG-ICG NPs in OVCAR8 cells (Figure S20); intracellular ROS generation following PCA1.5-PEG NP treatment in OVCAR3 cells (Figure S21); effect of hemoglobin, intralipid, and melanin on the optical properties of PCA1.5-PEG-ICG NPs (Figures S22-S24); TMP diameter uniformity and quantitative fluorescence imaging of PCA-PEG-ICG TMPs (Figures S25-S26); photobleaching experimental setup and NIR-I fluorescence imaging of PEG-ICG NP TMPs following white-light exposure (Figures S27-S28); tissue-stacking setup and tissue-stacking profiles using muscle, fat, and skin layers for PCA-PEG-ICG TMPs versus ICG TMPs (Figures S29-S32); 3D-bioprinted skin-mimicking constructs and 3D-bioprinted intraperitoneal tumor mimic with embedded TMPs (Figures S33-S34)

## ACKNOWLEDGMENT

I.S. acknowledges the Edward E. Whitacre Jr. College of Engineering, Texas Tech University and American Heart Association Career Development Award (26CDA1597763) for research support. All schematic figures were made in Biorender. Srivastava, I. (2026).

