## Supporting Information for "Architecture-Dependent Stability, Cellular Uptake, and Redox Modulation of Poly(p-Coumaric Acid) Hybrid Nanoparticles for Ovarian Carcinoma Intervention"

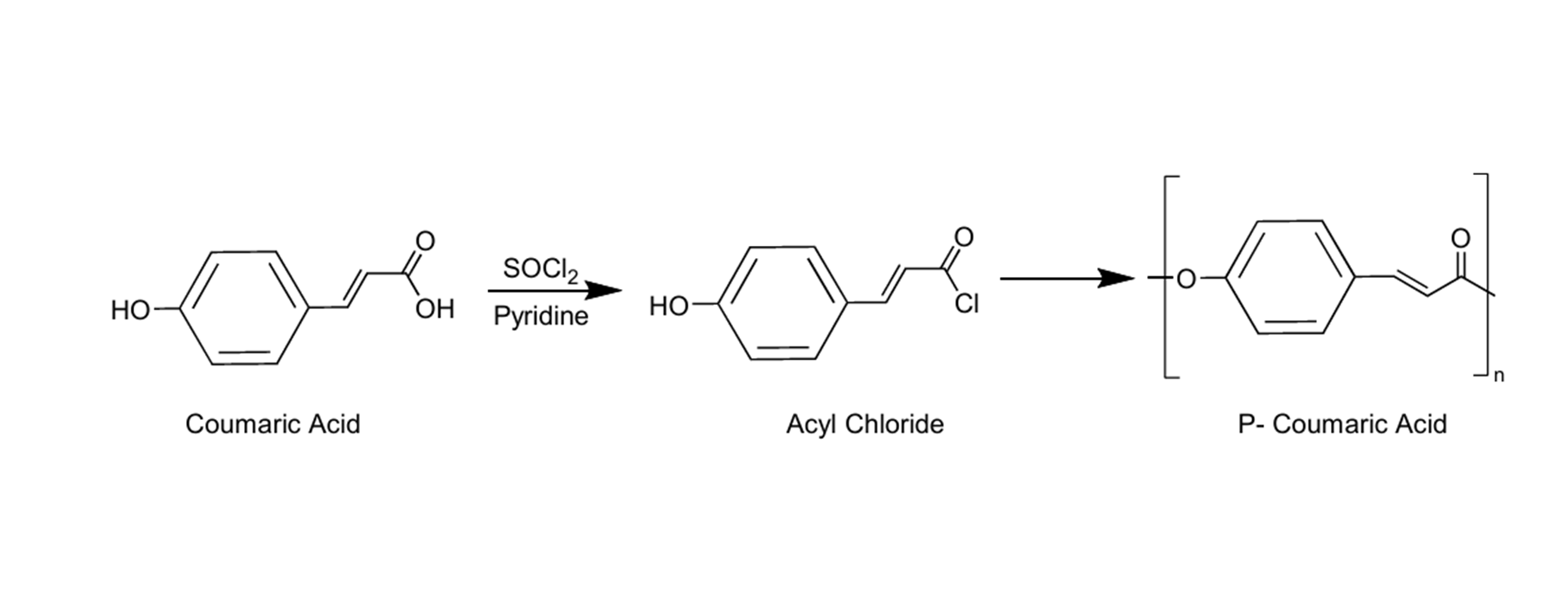

**Figure S1**. Schematic diagram of one-step polycondensation synthesis of PCA from CA.

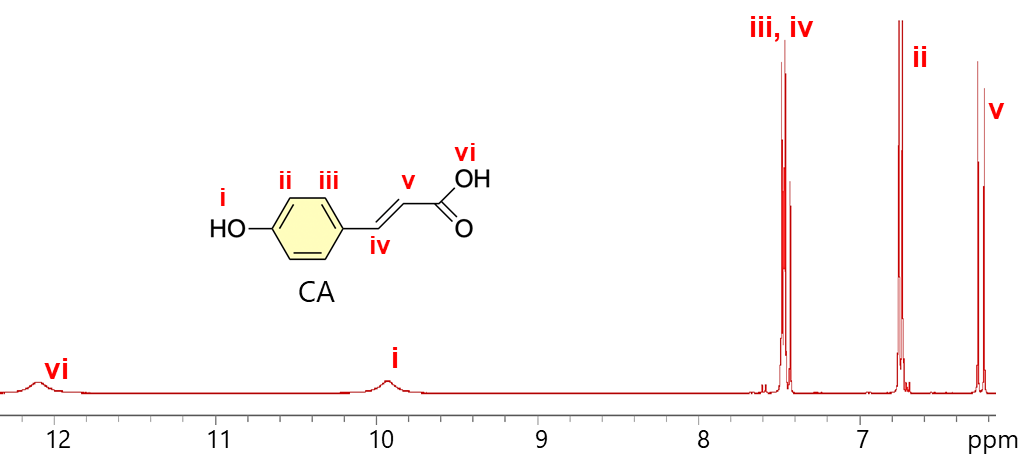

**Figure S2. ^1^H-NMR Characterization of CA.** ^1^H-NMR spectrum of CA.

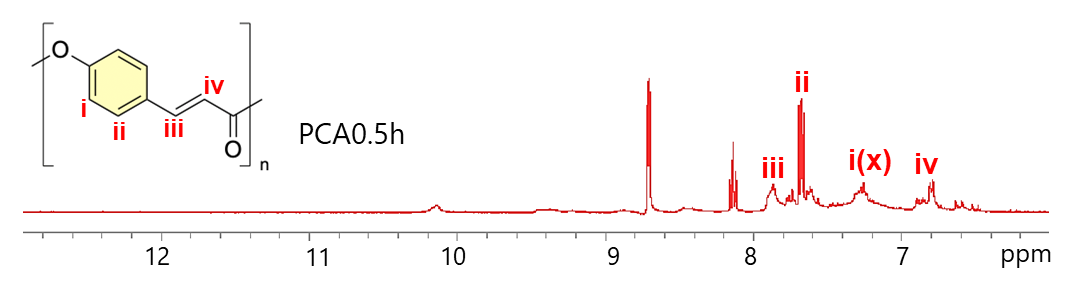

**Figure S3. ^1^H-NMR Characterization of PCA 0.5h.** ^1^H-NMR spectrum of PCA after 0.5h (PCA 0.5h) of reaction time.

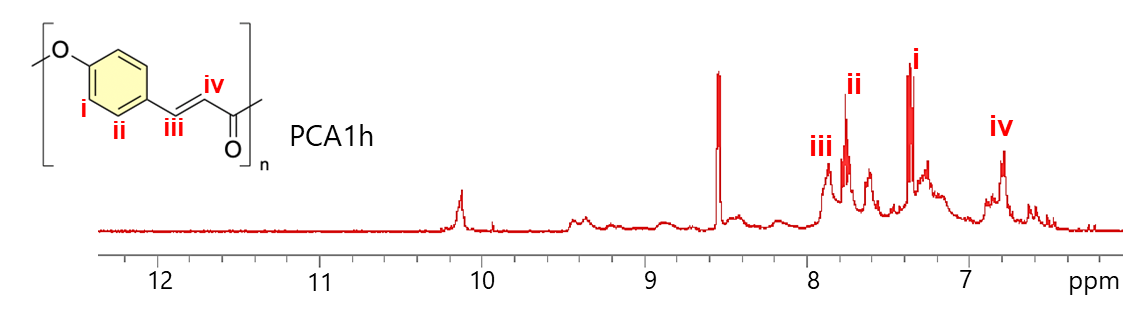

**Figure S4. ^1^H-NMR Characterization of PCA 1h.** ^1^H-NMR spectrum of PCA after 1h of reaction time (PCA 1h).

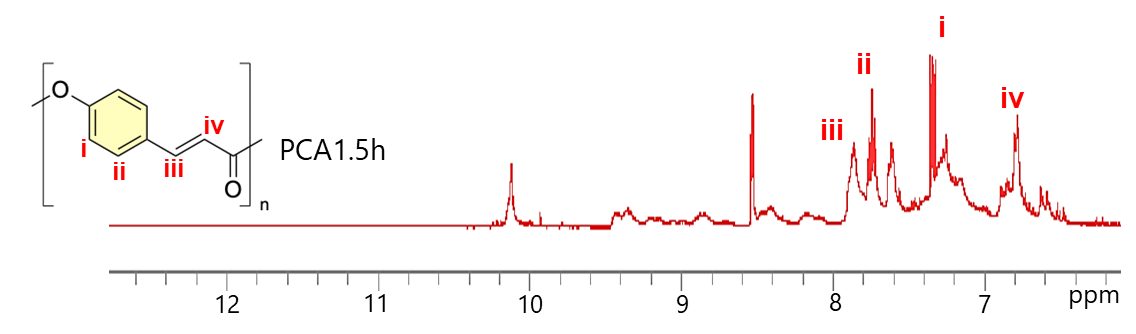

**Figure S5. ^1^H-NMR Characterization of PCA 1.5h.** ^1^H NMR spectrum of PCA after 1.5h of reaction time (PCA 1.5h).

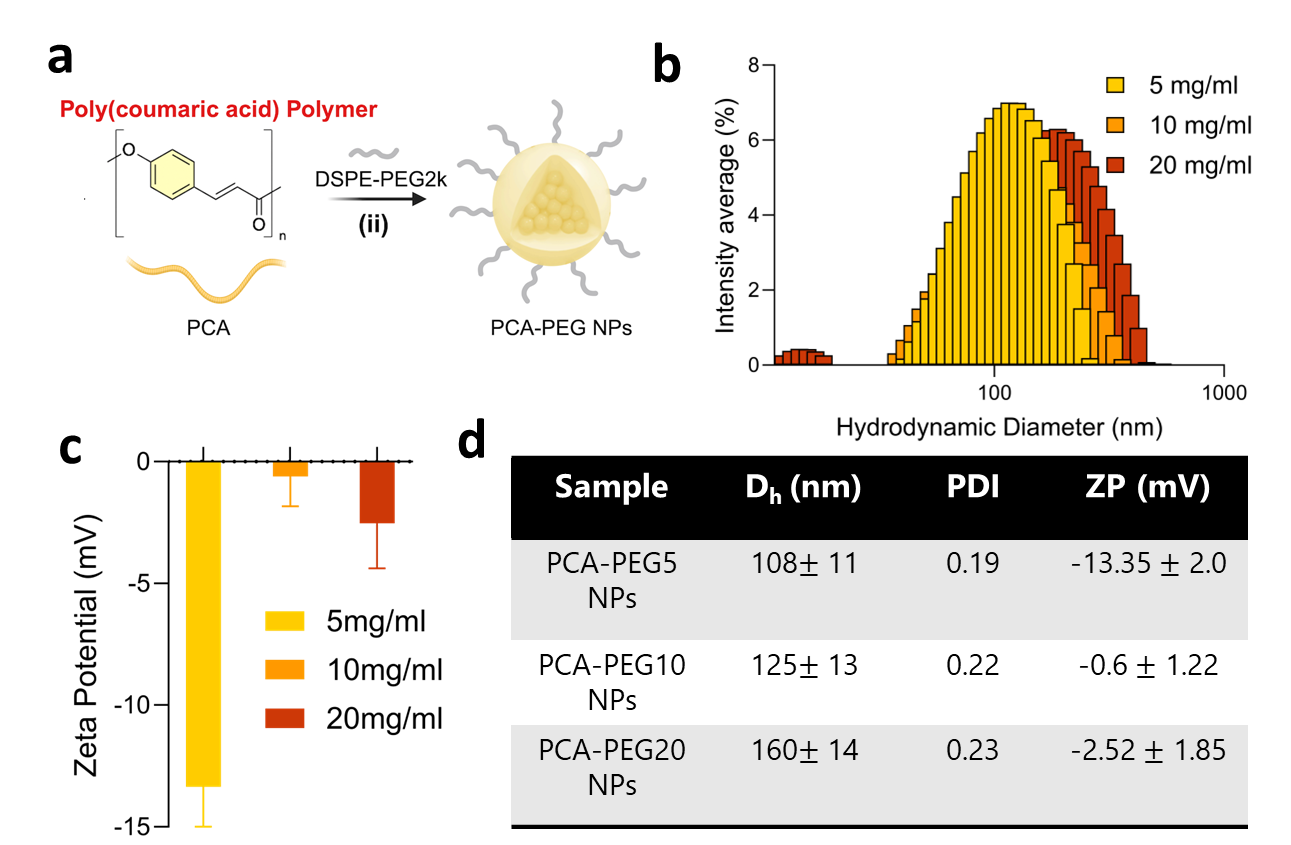

**Figure S6. Synthesis, optimization, and physicochemical characterization of PCA1.5-PEG NPs. (**a) Schematic overview of PCA-PEG NPs synthesis using PCA polymer (b) Hydrodynamic diameter intensity graph of PCA1.5h-PEG5 NPs, PCA1.5h-PEG10 NPs and PCA1.5h-PEG20 NPs (c) ζ-potential graph of PCA1.5h-PEG5 NPs, PCA1.5h-PEG10 NPs and PCA1.5h-PEG20 NPs (d) Hydrodynamic diameter, PDI, and ζ-potential values of PCA-PEG5 NPs, PCA-PEG10 NPs and PCA-PEG20 NPs.

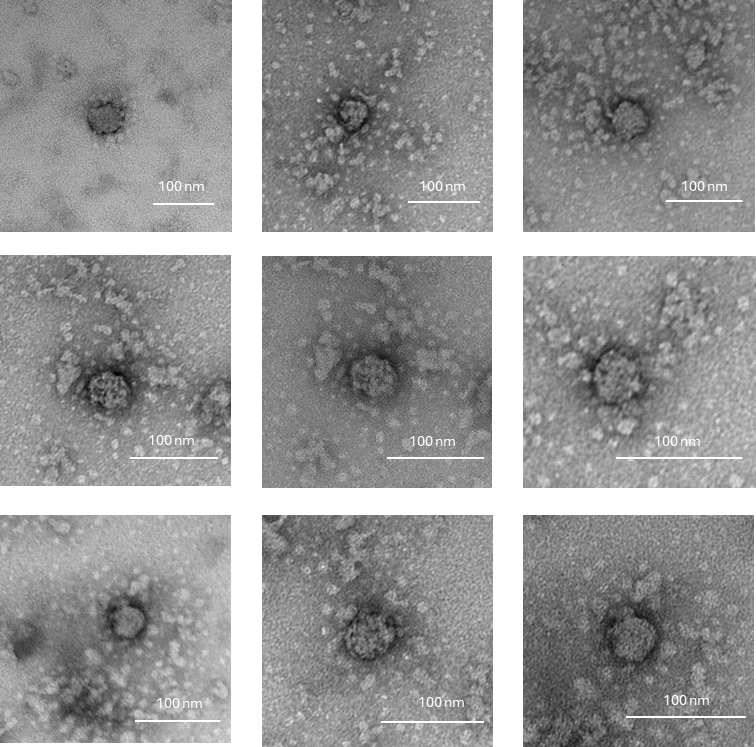

**Figure S7.** Negative-stained TEM images of PCA0.5h-PEG NPs.

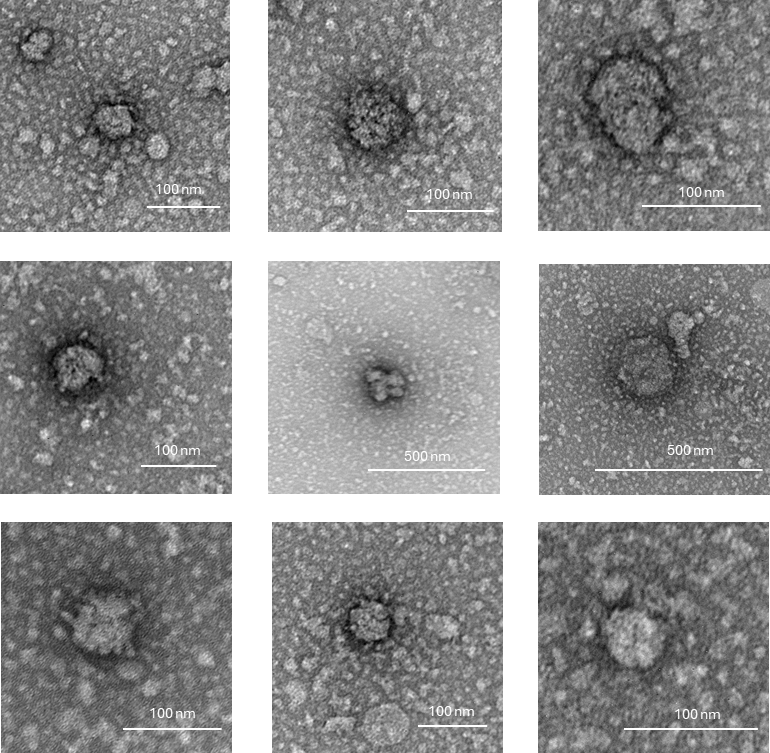

**Figure S8.** Negative-stained TEM images of PCA1h-PEG NPs.

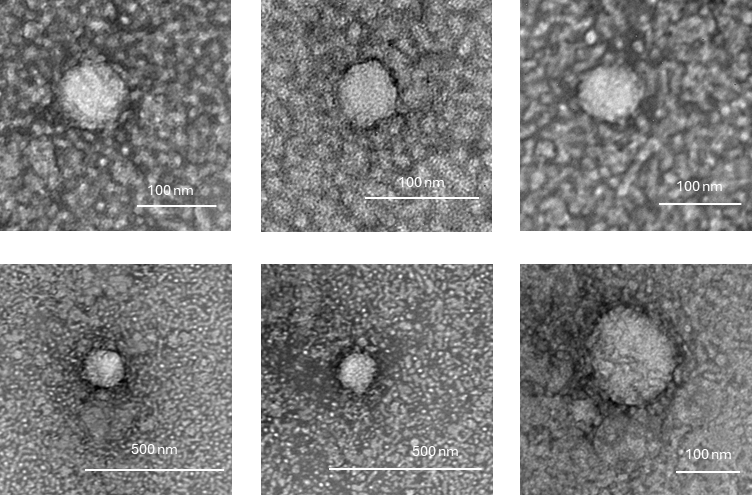

**Figure S9.** Negative-stained TEM images of PCA1.5h-PEG NPs

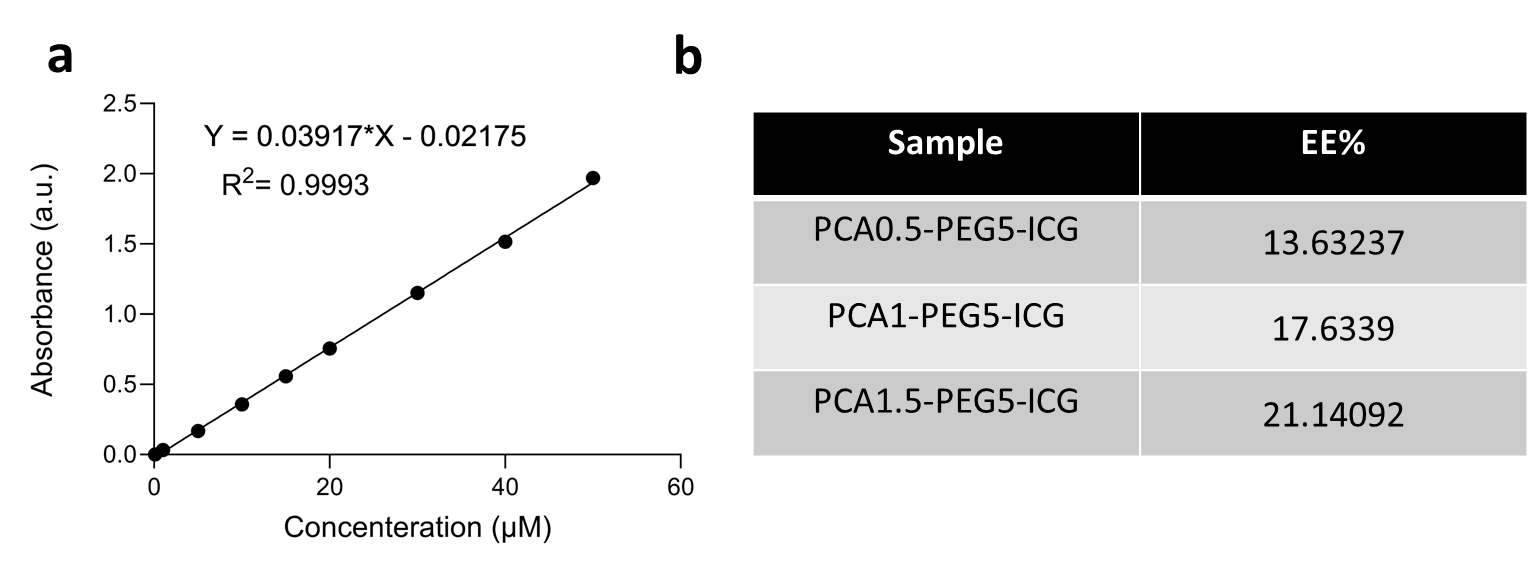
**Figure S10. Encapsulation Efficiency of ICG in PCA-PEG-ICG NPs.** (a) Calibration curve showing variation of ICG concentration versus absorbance at 702 nm. (b) The encapsulation efficiency (EE) of ICG in PCA0.5-PEG-ICG, PCA1-PEG-ICG and PCA1.5-PEG-ICG NPs.

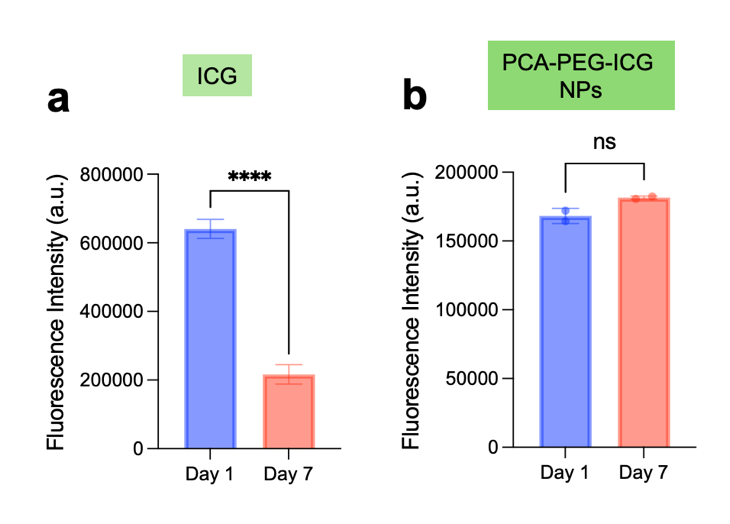

**Figure S11. Quantitative fluorescence stability of free ICG and PCA1.5-PEG-ICG NPs over 7 days.** Fluorescence intensity of (a) free ICG and (b) PCA1.5-PEG-ICG NPs measured on Day 1 and Day 7. Free ICG showed a significant decrease in fluorescence intensity after 7 days, whereas PCA1.5-PEG-ICG NPs maintained fluorescence intensity over the same period. Data are presented as mean ± SD, n = 3. Statistical comparisons between Day 1 and Day 7 were performed using a two-tailed paired t-test. ns, not significant; ****p < 0.0001.

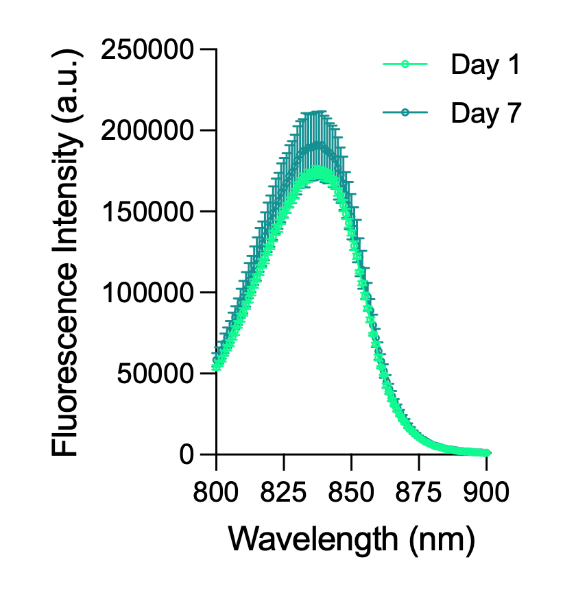

**Figure S12. Time-dependent fluorescence stability of PEG-ICG NPs.** Fluorescence intensity of PEG-ICG NPs was measured on day 1 and day 7 to evaluate nanoparticle-associated ICG signal retention over time. Data are presented as mean ± SD, n = 3.

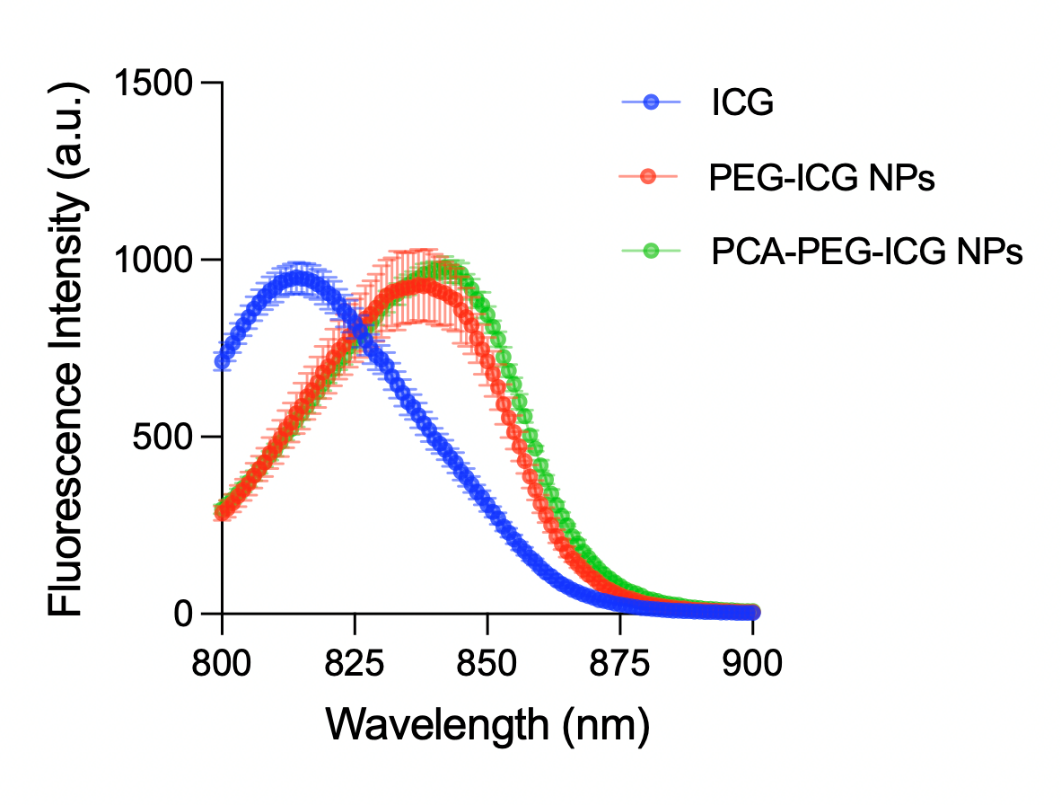

**Figure S13. Comparative fluorescence stability of free and nanoparticle-associated ICG formulations.** Fluorescence intensity of free ICG, PEG-ICG NPs, and PCA1.5-PEG-ICG NPs was compared on day 7 to assess the effect of nanoparticle formulation on ICG signal stability. Data are presented as mean ± SD, n = 3.

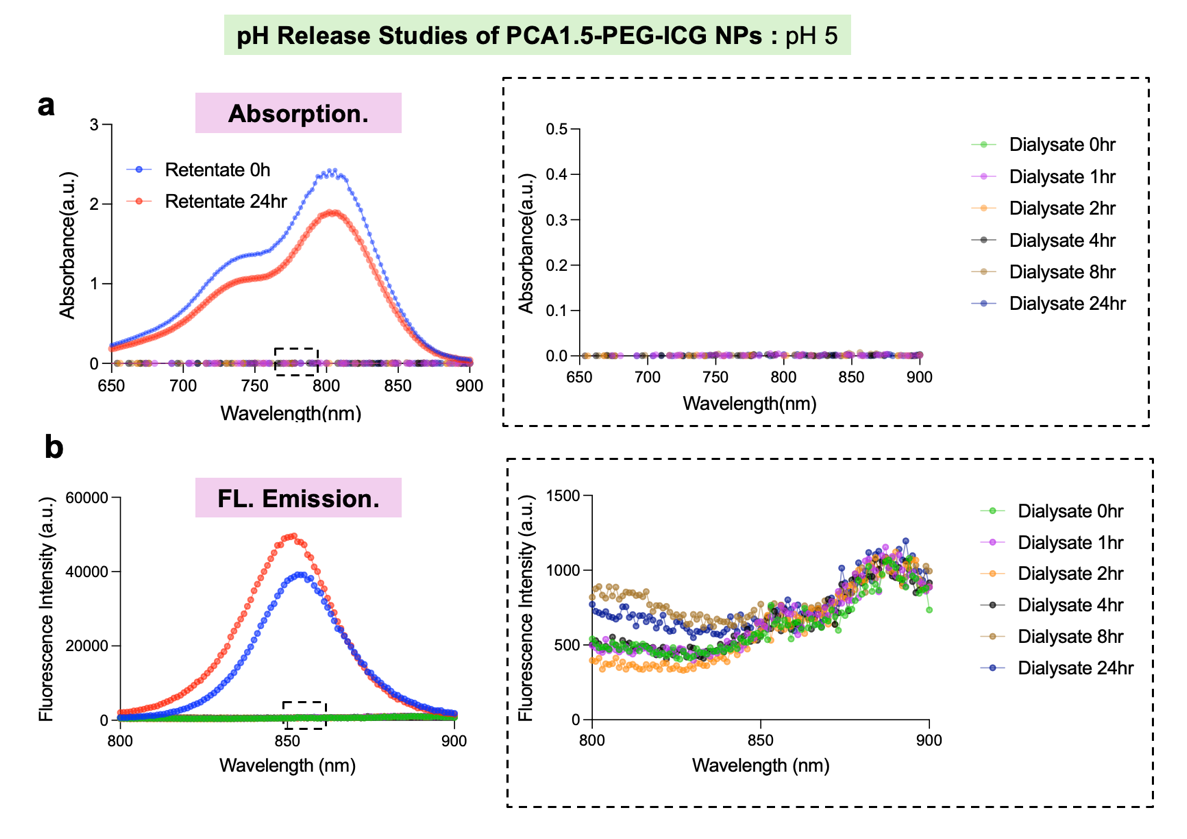

**Figure S14. *In vitro* ICG release assessment of PCA1.5-PEG-ICG NPs under acidic pH conditions.** PCA1.5-PEG-ICG NPs were dialyzed at pH 5.0 for 24 h to evaluate ICG retention and release under lysosomal-mimicking acidic conditions. (a) UV–Vis absorbance spectra of the nanoparticle retentate at 0 and 24 h and dialysate fractions collected over time. (b) Fluorescence emission spectra of the corresponding retentate and dialysate fractions. The retentate retained the characteristic optical signal of encapsulated ICG, whereas the dialysate fractions showed minimal absorbance and weak fluorescence signal, indicating limited ICG release from PCA1.5-PEG-ICG NPs under acidic conditions over 24 h.

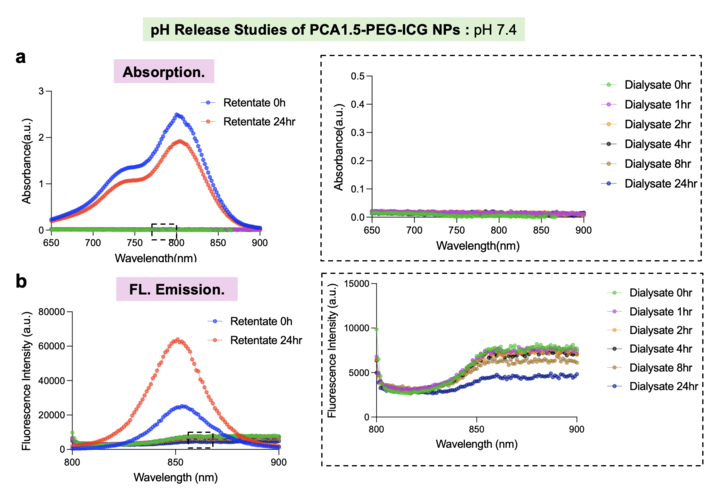

**Figure S15. *In vitro* ICG release assessment of PCA1.5-PEG-ICG NPs under physiological pH conditions.** PCA1.5-PEG-ICG NPs were dialyzed at pH 7.4 for 24 h to evaluate ICG retention and release under physiological pH conditions. (a) UV–Vis absorbance spectra of the nanoparticle retentate at 0 and 24 h and dialysate fractions collected over time. (b) Fluorescence emission spectra of the corresponding retentate and dialysate fractions. The retentate retained the characteristic optical signal of encapsulated ICG, whereas the dialysate fractions showed minimal absorbance and low fluorescence signal relative to the retentate, indicating limited detectable ICG release from PCA1.5-PEG-ICG NPs under physiological conditions over 24 h.

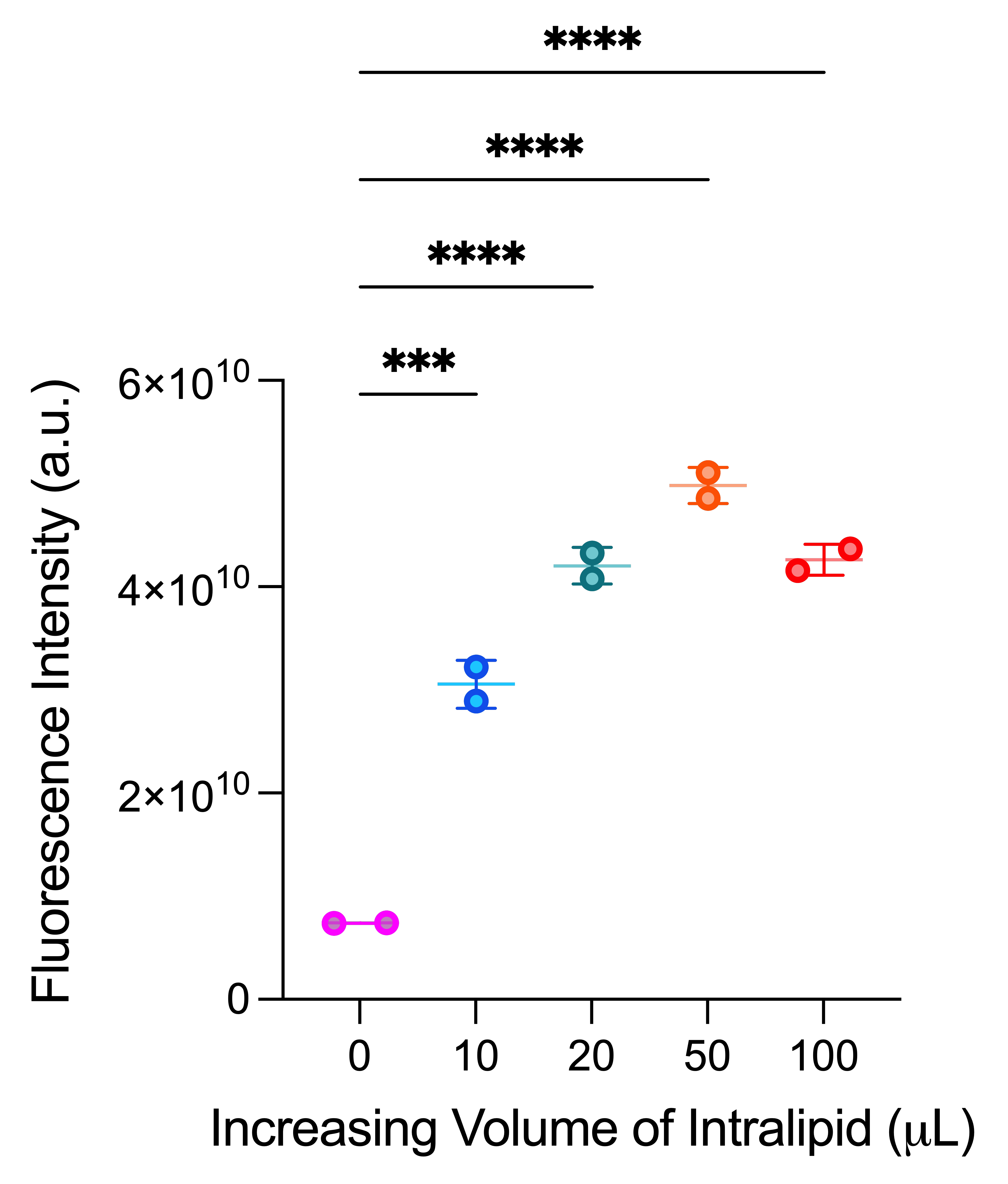

**Figure S16.** Impact of increasing Intralipid concentration on the optical performance of PCA-PEG-ICG NP. NIR-I fluorescence image was acquired using IVIS imaging system at an excitation wavelength of 745 nm and emission wavelength of 840 nm.

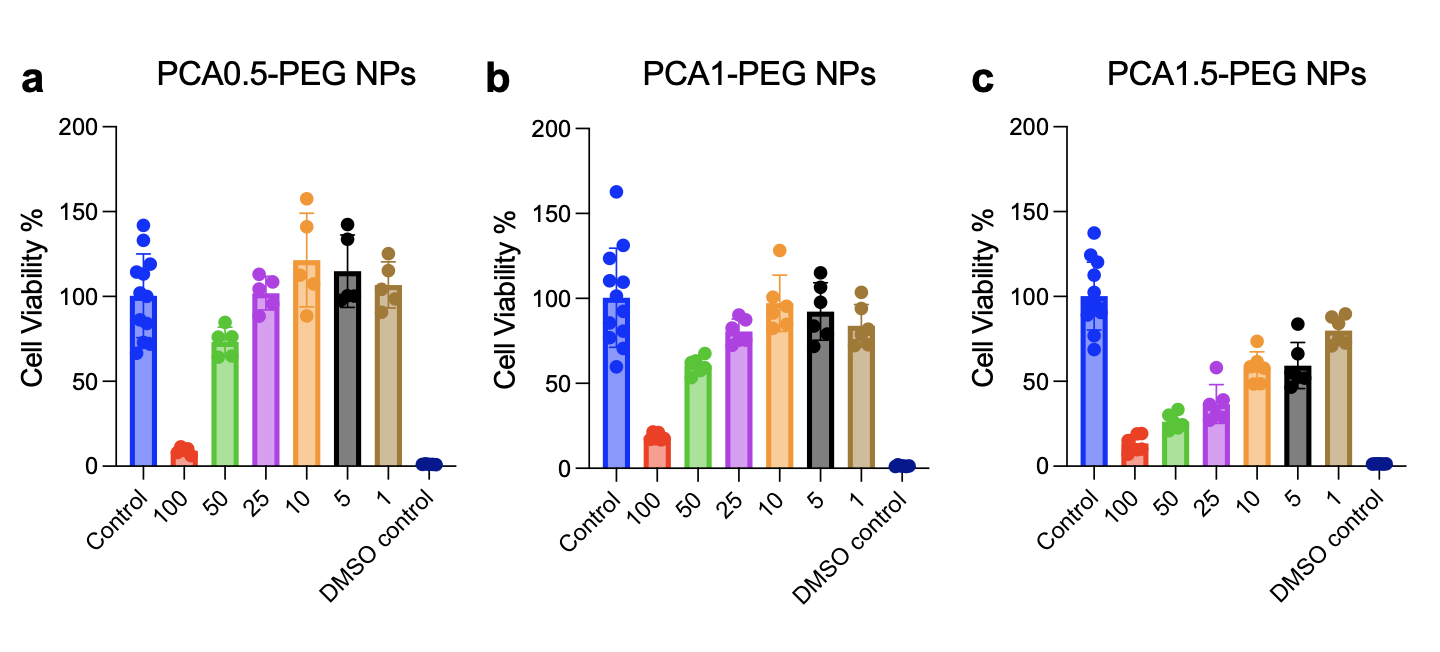
**Figure S17. *In vitro* evaluation of cytotoxicity of PCA-PEG NPs in OVCAR3 cells.** Cell viability of OVCAR3 following treatment with **(a)** PCA0.5-PEG NPs, **(b)** PCA1-PEG NPs, and **(c)** PCA1.5-PEG NPs at the indicated concentrations. Data are presented as mean ± SD.

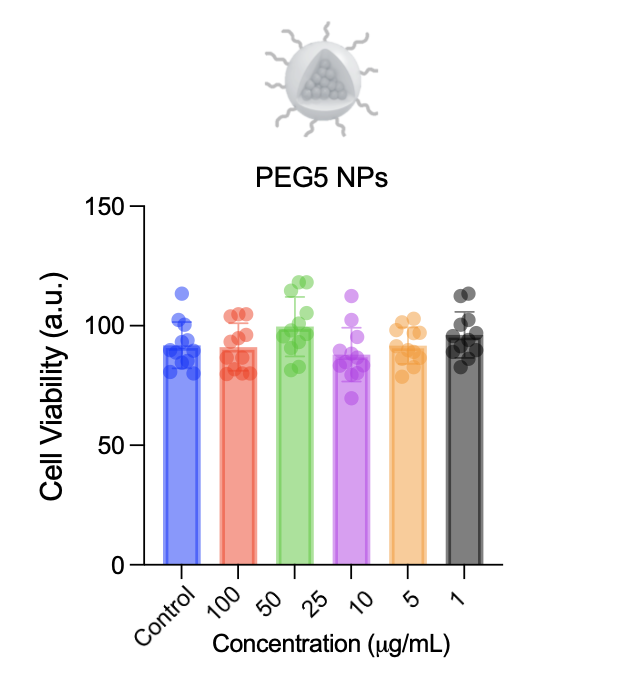

**Figure S18.** Cell viability of OVCAR8 following treatment with PEG5 NPs (with no PCA).

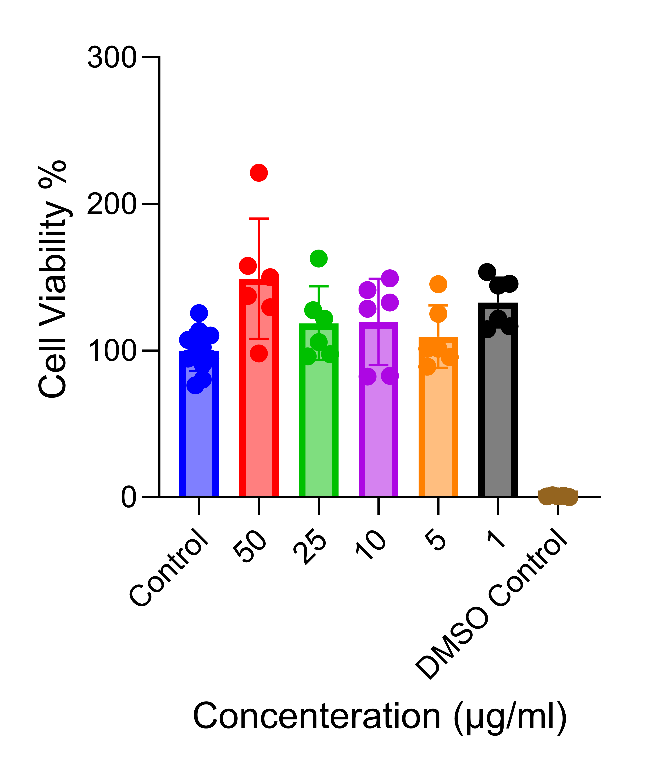

**Figure S19. *In vitro* evaluation of cytotoxicity of PCA-PEG NPs in HUVECs.** Cell viability of HUVECs following treatment with PCA1.5-PEG NPs at the indicated concentrations. Data are presented as mean ± SD.

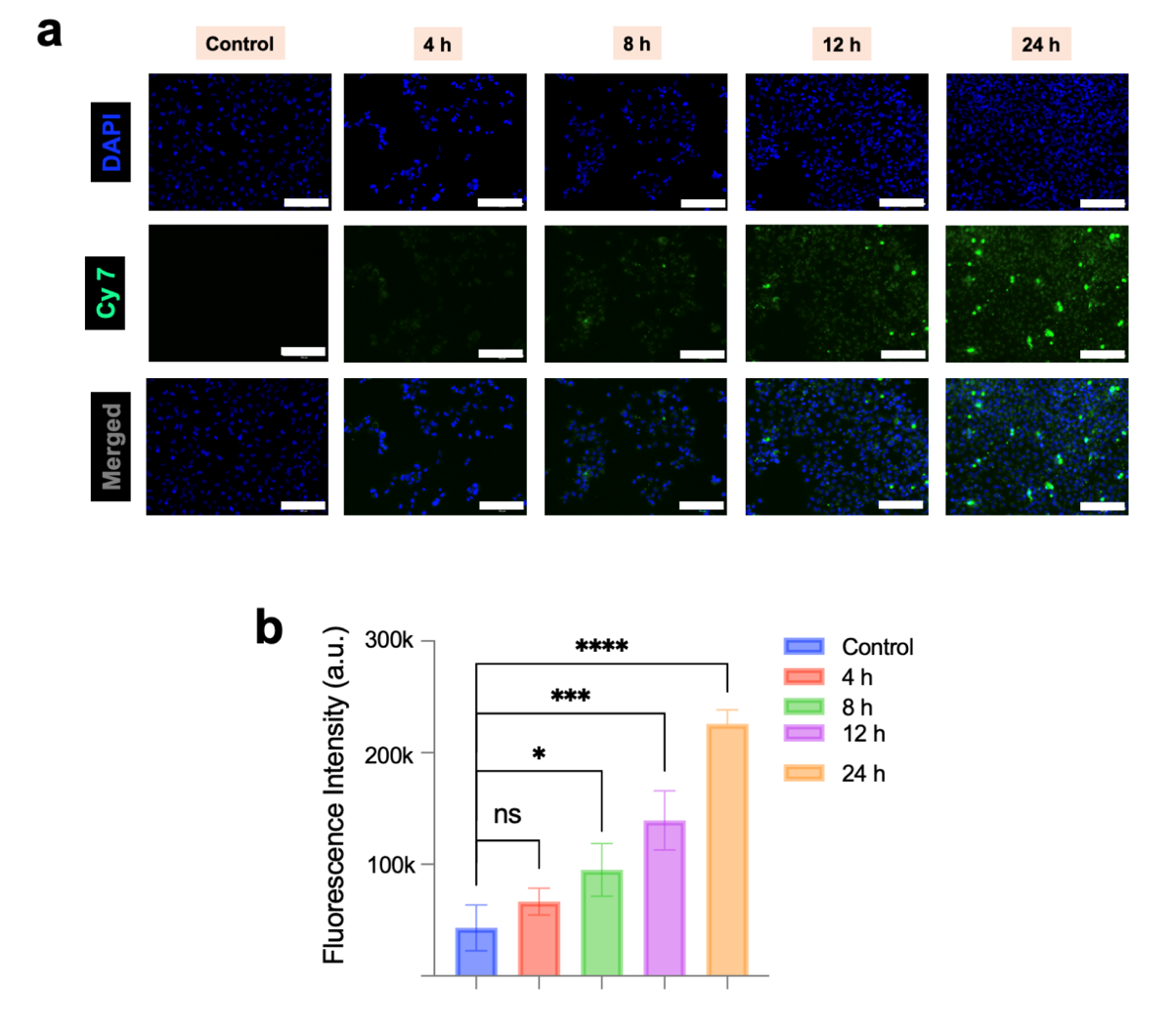

**Figure S20. Time-dependent cellular uptake of PCA1.5-PEG-ICG NPs in OVCAR8 cells.**
(a) Representative fluorescence microscopy images of OVCAR8 cells following incubation with PCA1.5-PEG-ICG NPs for 4, 8, 12, and 24 h, with untreated cells serving as controls. Nuclei are shown in blue (DAPI), and ICG-associated fluorescence is shown in green (Cy7 channel). Scale bars = *150 μm*. (b) Quantification of cellular fluorescence intensity following treatment with PCA1.5-PEG-ICG NPs at the indicated time points compared with untreated controls. Data are presented as mean ± SD, n = 4. Statistical comparisons were performed using one-way ANOVA followed by Dunnett’s multiple-comparisons test versus untreated control. ns, not significant; *p < 0.05; ***p < 0.001; ****p < 0.0001.

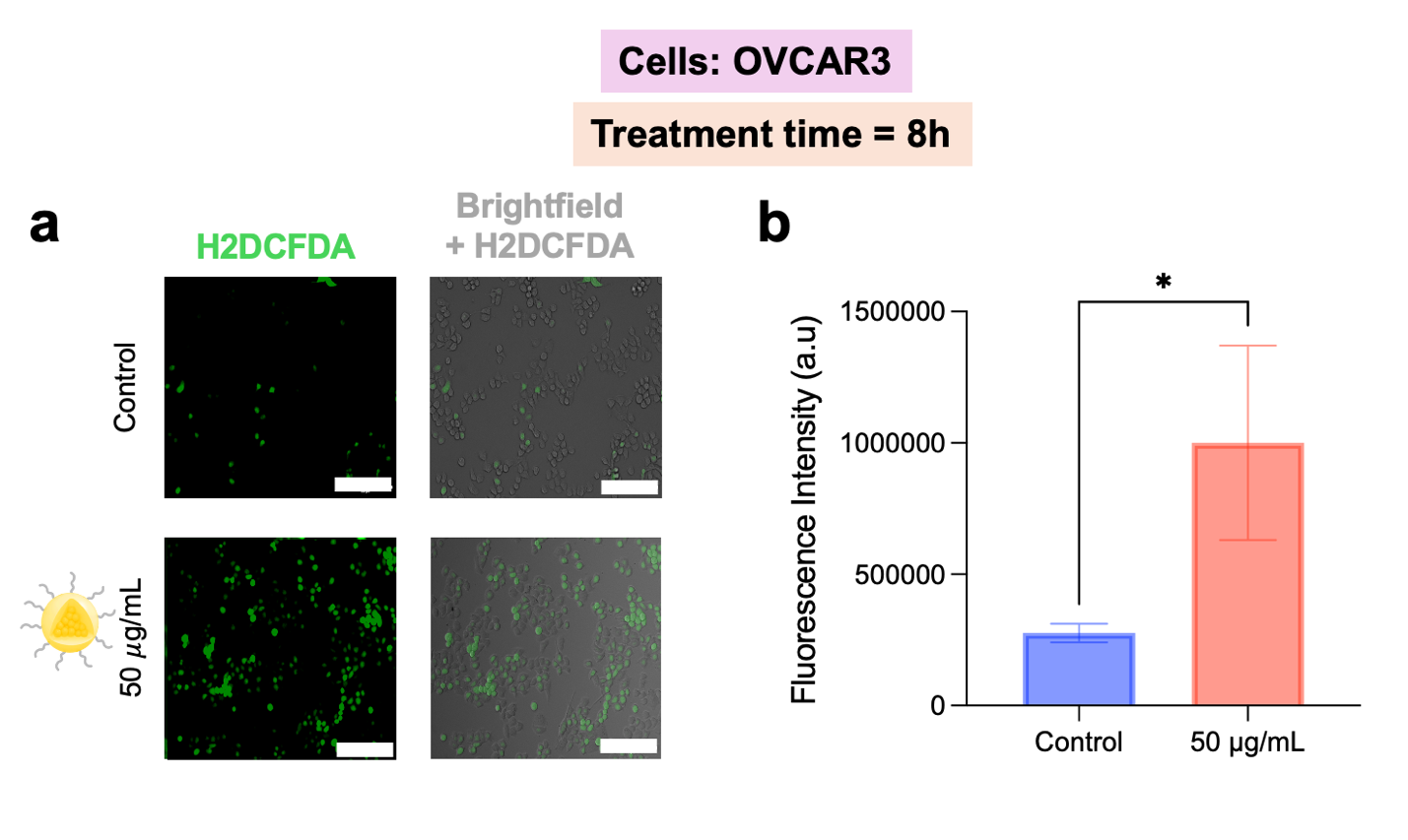

**Figure S21. *In vitro* assessment of intracellular ROS generation following PCA1.5-PEG NP treatment in OVCAR 3 cells.** **(a)** Representative fluorescence and brightfield-overlay images of OVCAR 3 cells stained with H2DCFDA following treatment with PCA1.5-PEG NPs at 50 µg/mL for 8 h, with untreated cells serving as controls. **(b)** Quantification of H2DCFDA fluorescence intensity in control and PCA1.5-PEG NP-treated OVCAR3 cells. Data are presented as mean ± SD. Statistical significance is indicated as p < 0.05. Scale bar is 150 μm.

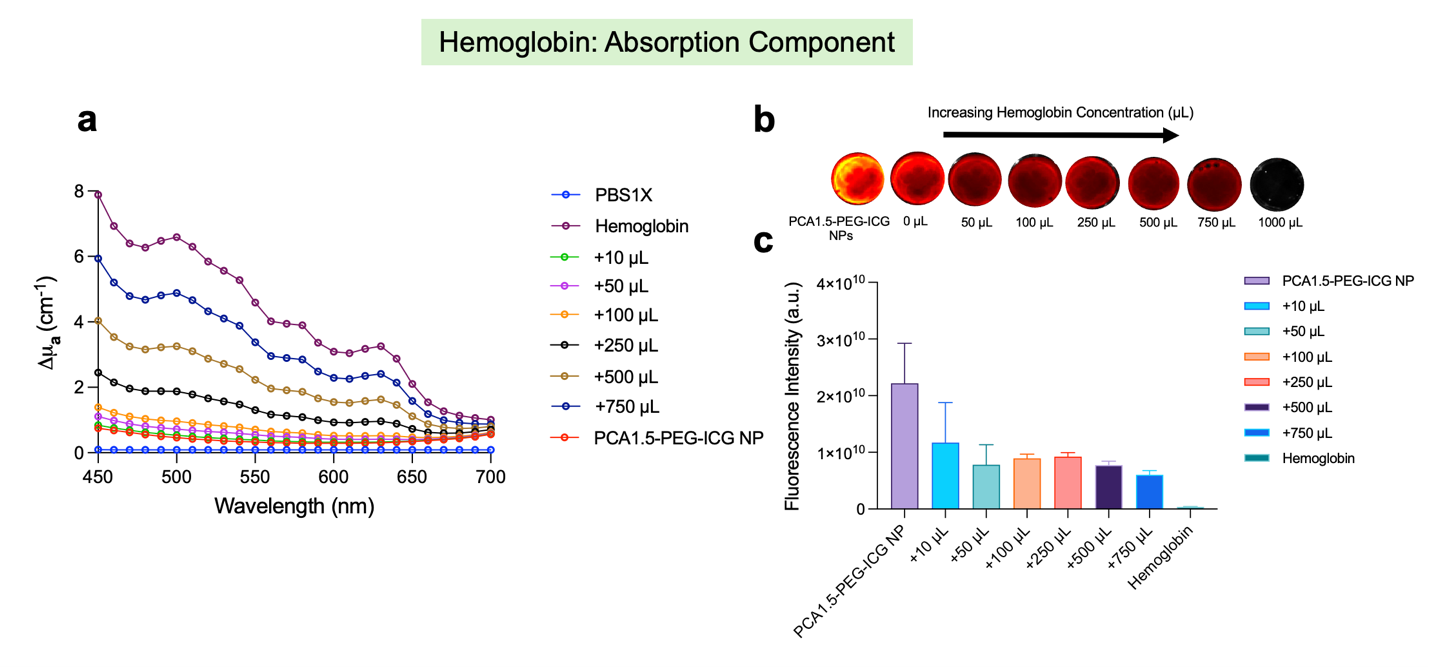

**Figure S22. Effect of hemoglobin concentration on the optical properties of PCA1.5-PEG-ICG NPs.** **(a)** Absorption coefficient change (Δµ_a_) of PCA1.5-PEG-ICG NPs in the presence of increasing hemoglobin concentrations. **(b)** Representative fluorescence images of PCA1.5-PEG-ICG NPs with increasing hemoglobin content ranging from 10 to 1000 µL. **(c)** Quantification of fluorescence intensity as a function of hemoglobin content. Data are presented as mean ± SD.

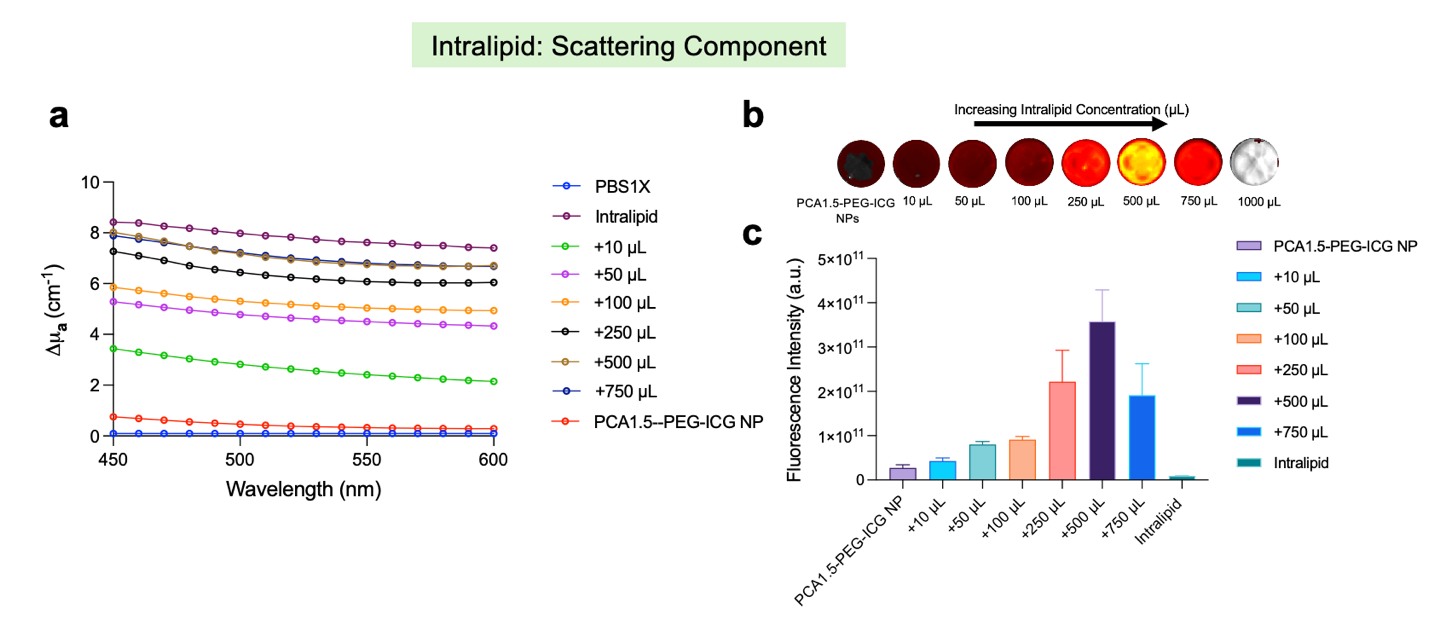

**Figure S23. Effect of Intralipid as a scattering component on PCA1.5-PEG-ICG NP fluorescence.** **(a)** Absorption coefficient change (Δµ_a_) of PCA1.5-PEG-ICG NPs measured with increasing Intralipid concentrations. **(b)** Representative fluorescence images of PCA1.5-PEG-ICG NPs containing increasing amounts of Intralipid from 10 to 1000 µL. **(c)** Quantification of fluorescence intensity as a function of Intralipid content. Data are presented as mean ± SD.

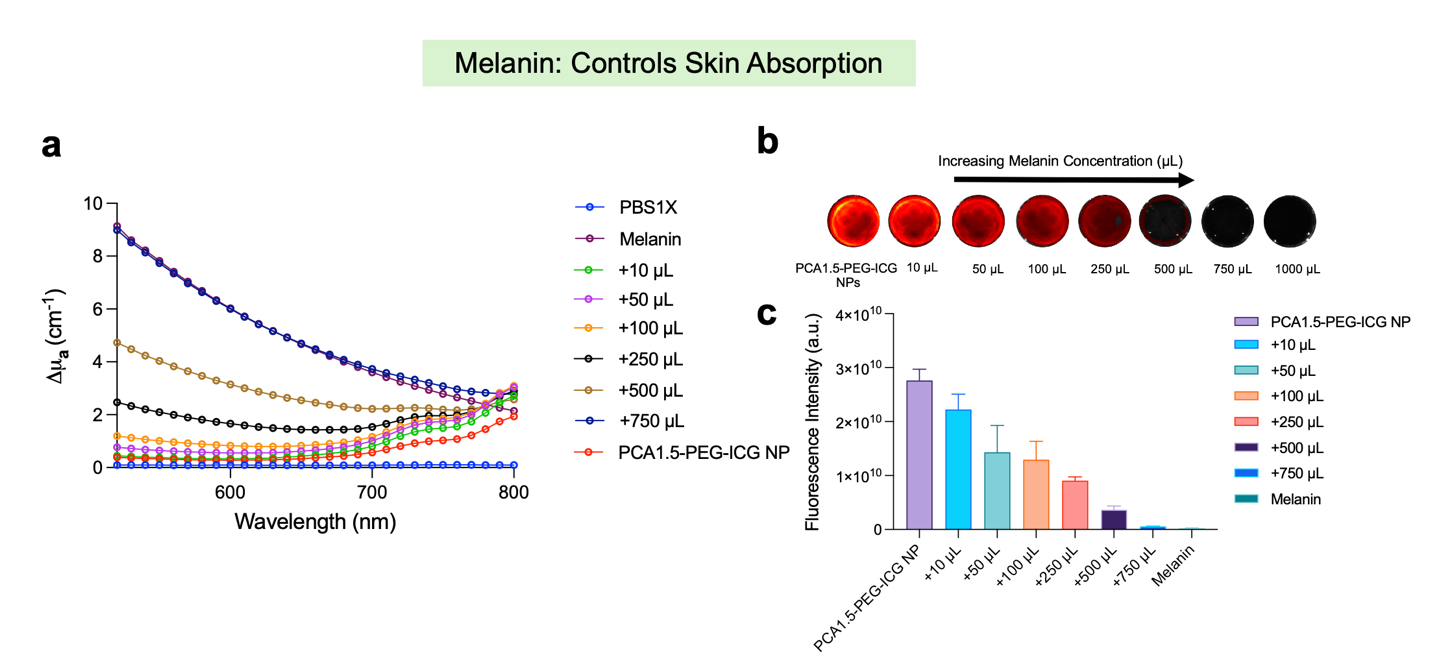

**Figure S24. Effect of melanin as a skin absorption component on PCA1.5-PEG-ICG NP fluorescence.** **(a)**Absorption coefficient change (Δµ_a_) of PCA1.5-PEG-ICG NPs measured with increasing melanin concentrations. **(b)** Representative fluorescence images of PCA1.5-PEG-ICG NPs containing increasing amounts of melanin from 10 to 1000 µL. **(c)** Quantification of fluorescence intensity as a function of melanin content. Data are presented as mean ± SD.

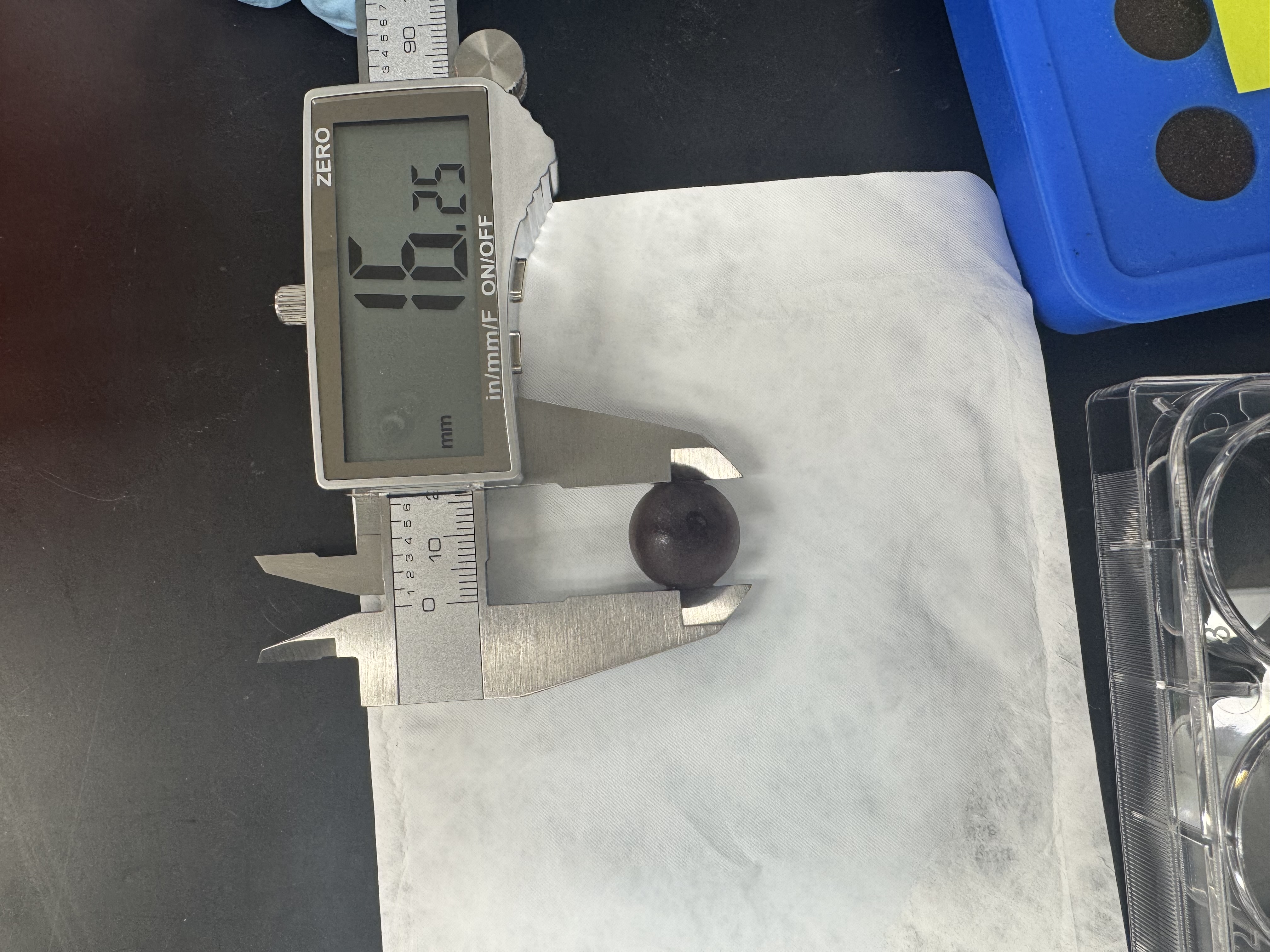

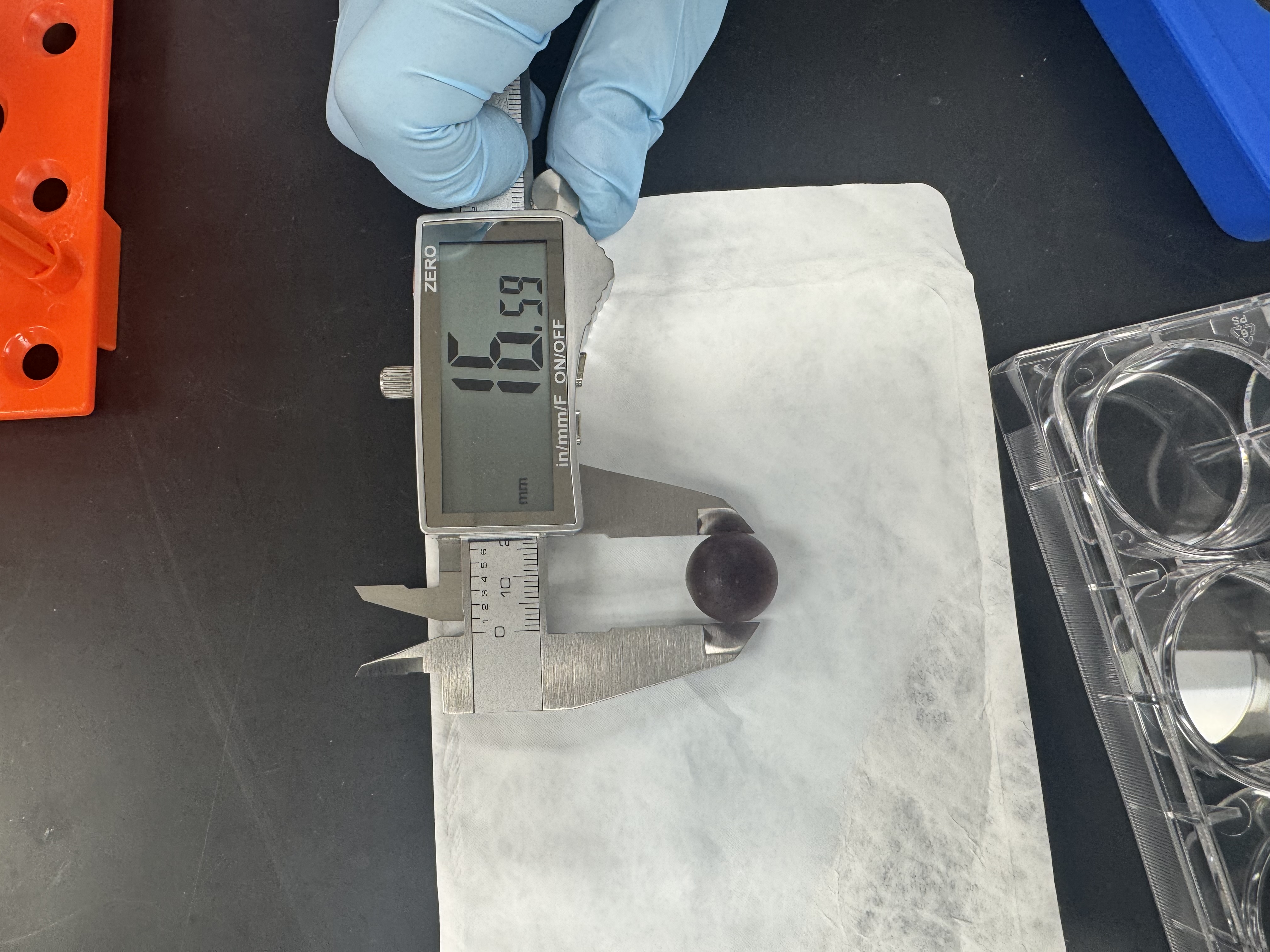

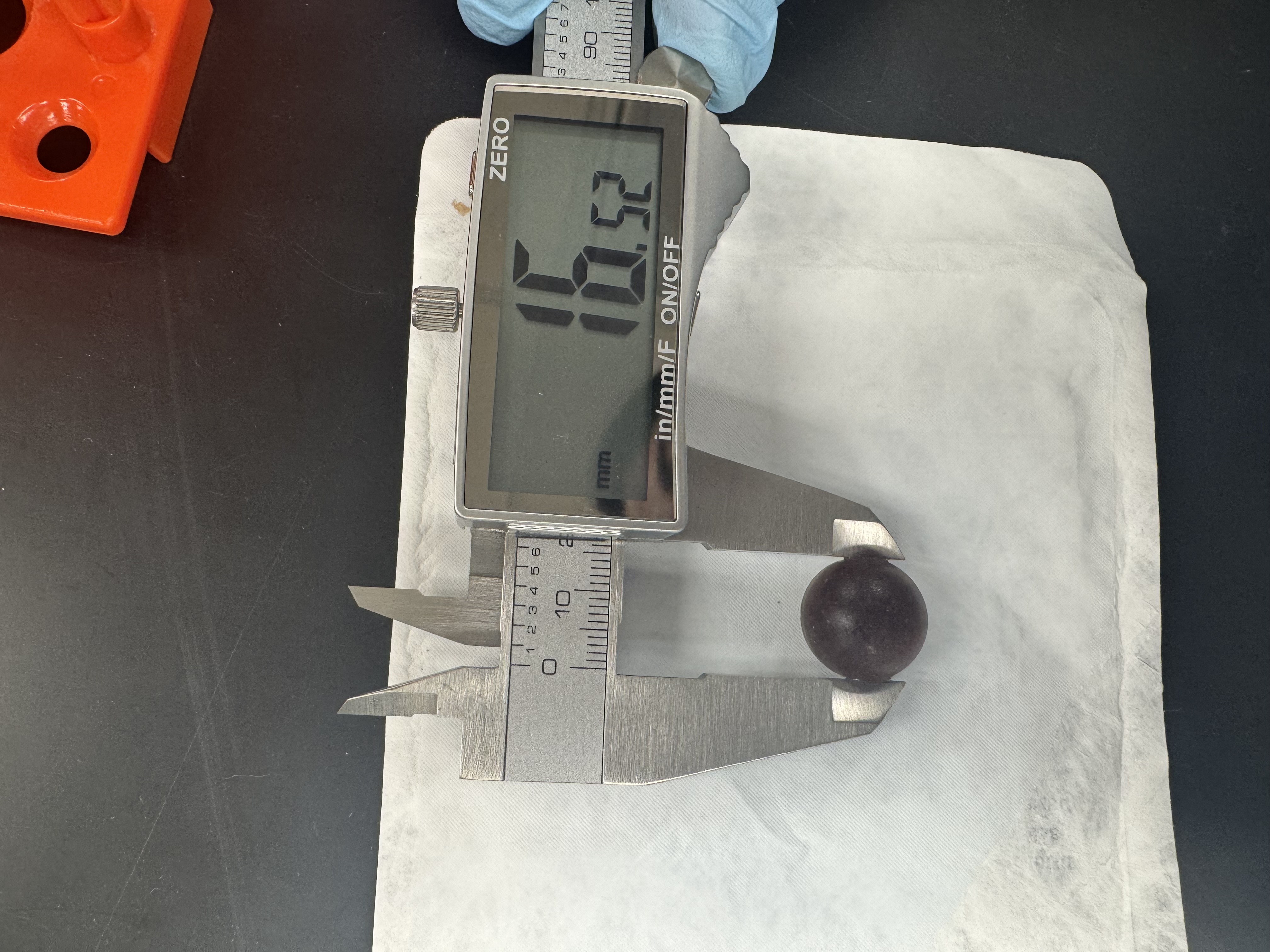

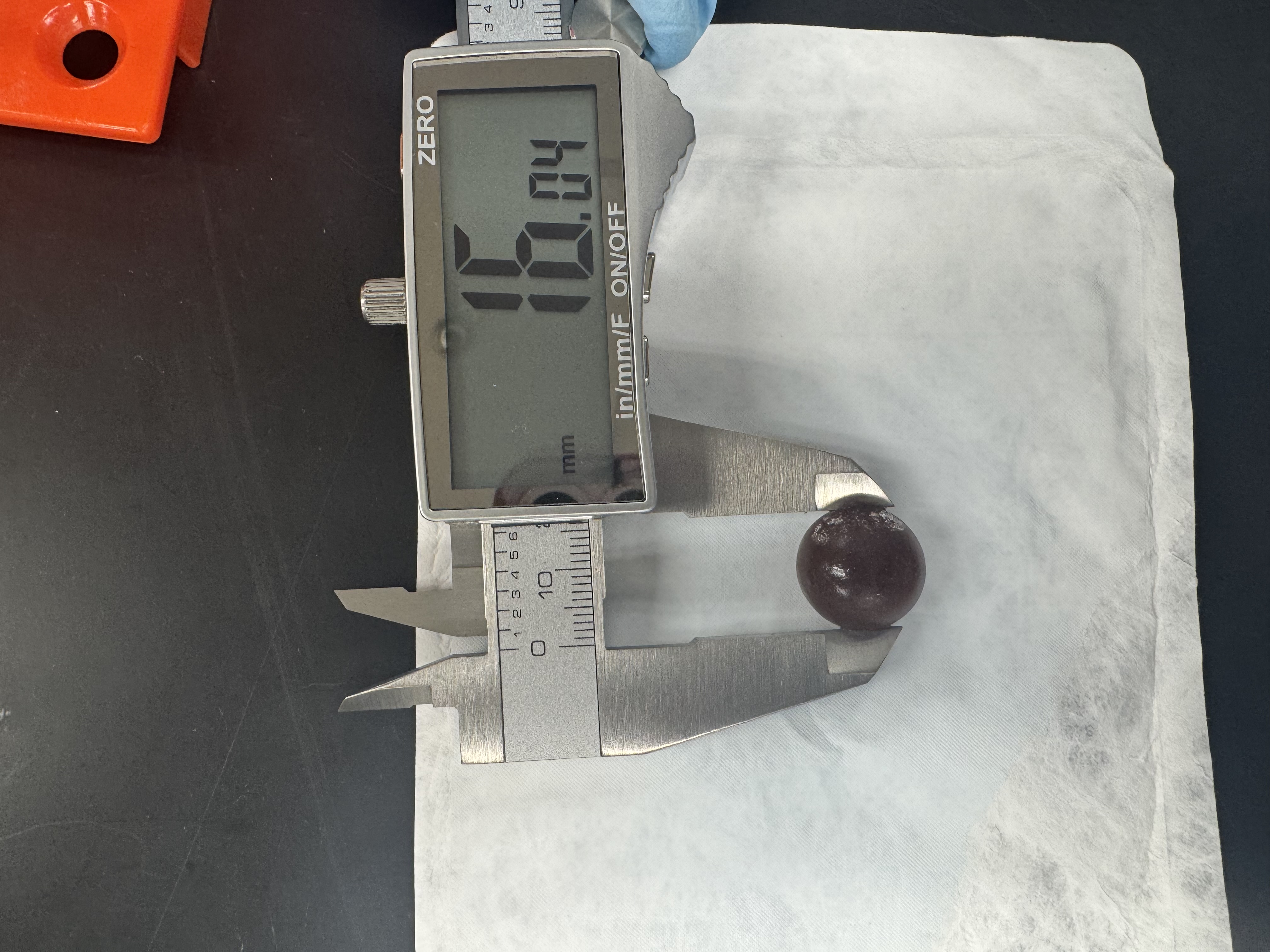

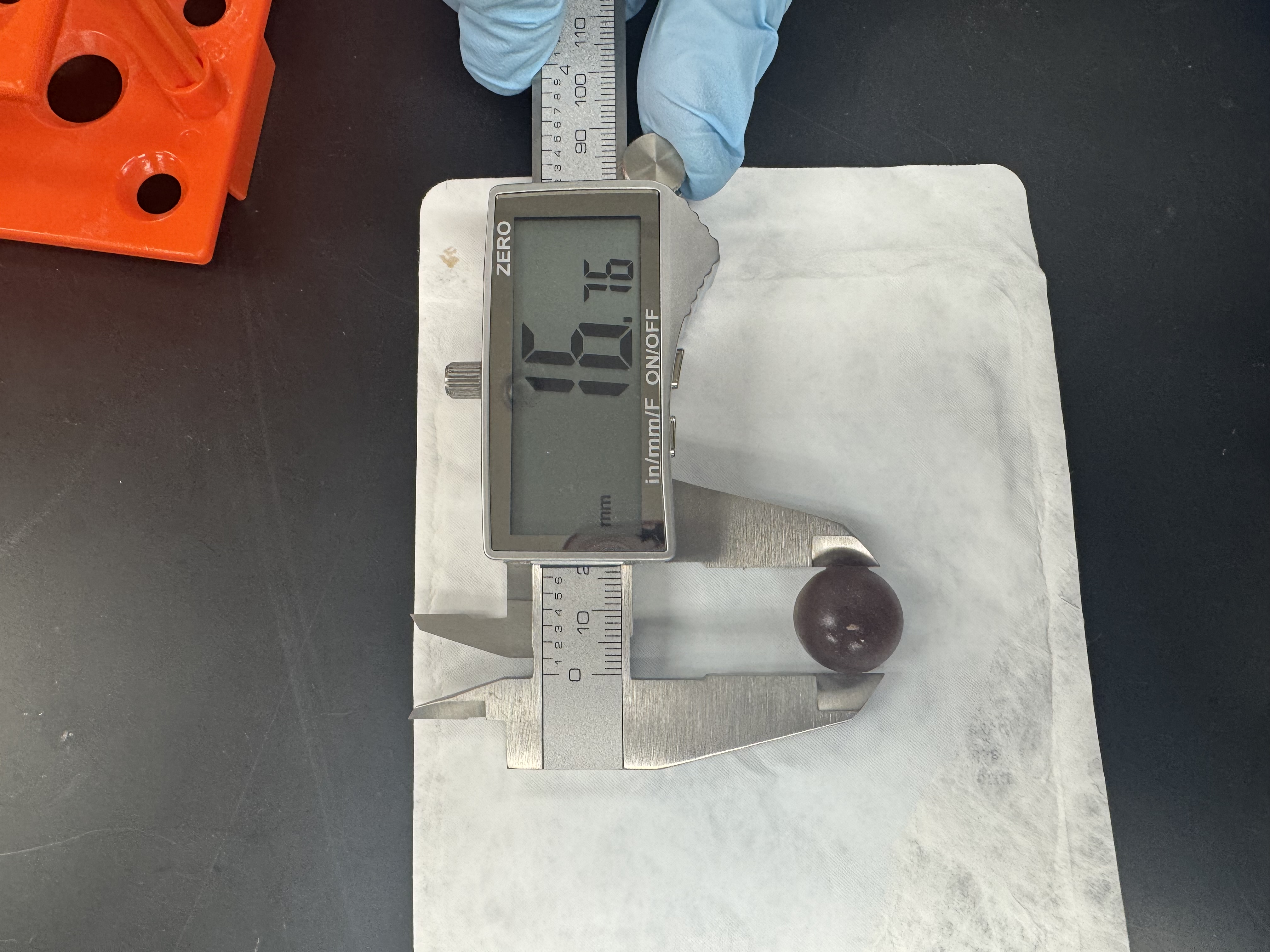

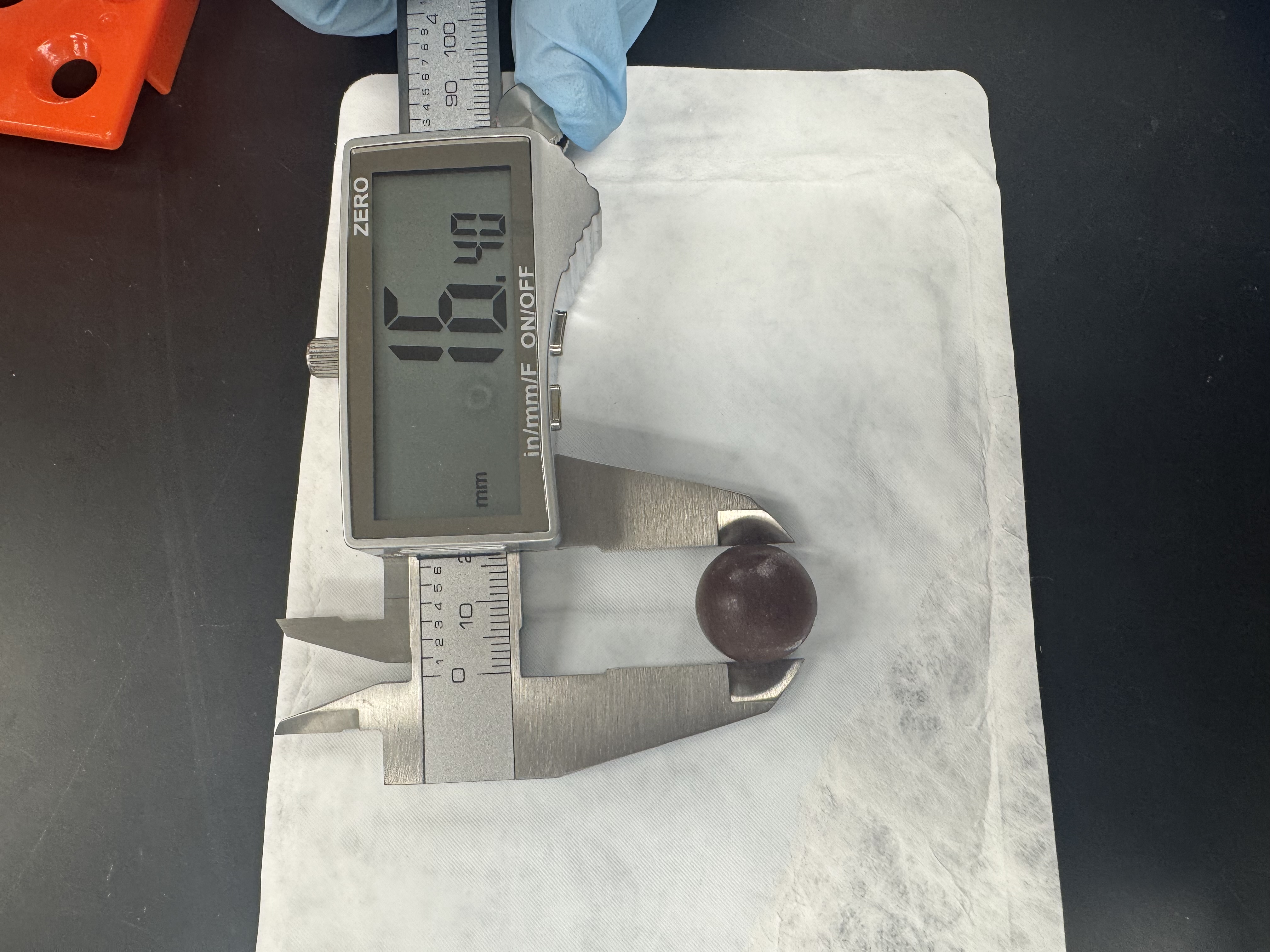

PBS1X

0.31 µg/mL

0.95 µg/mL

2.2 µg/mL

4.9 µg/mL

10 µg/mL

**Figure S25.** Representative images of the diameter of the prepared TMPs using digital calipers to show uniformity in our TMP preparation procedure. Concentration dilution was conducted by varying the concentration of ICG in PCA1.5h-PEG-ICG NPs and denoted on top of each figure.

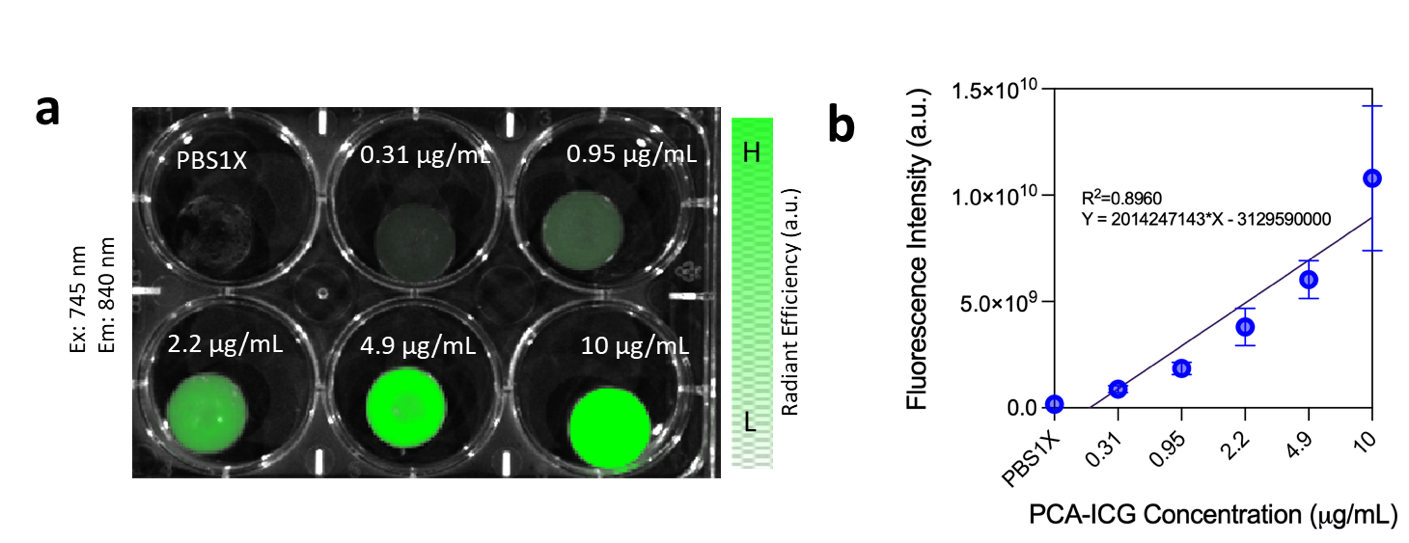

**Figure S26. Quantitative Fluorescence Imaging of PCA-PEG-ICG TMPs.** (a) NIR-I fluorescence image of PCA1.5h-PEG-ICG NPs *via* IVIS at varying concentrations. (b) The corresponding fluorescence intensities were acquired, and a linear signal decay trend was observed in serially diluted phantoms. Concentration dilution was conducted by varying the concentration of ICG in PCA1.5h-PEG-ICG NPs and denoted on top of each figure.

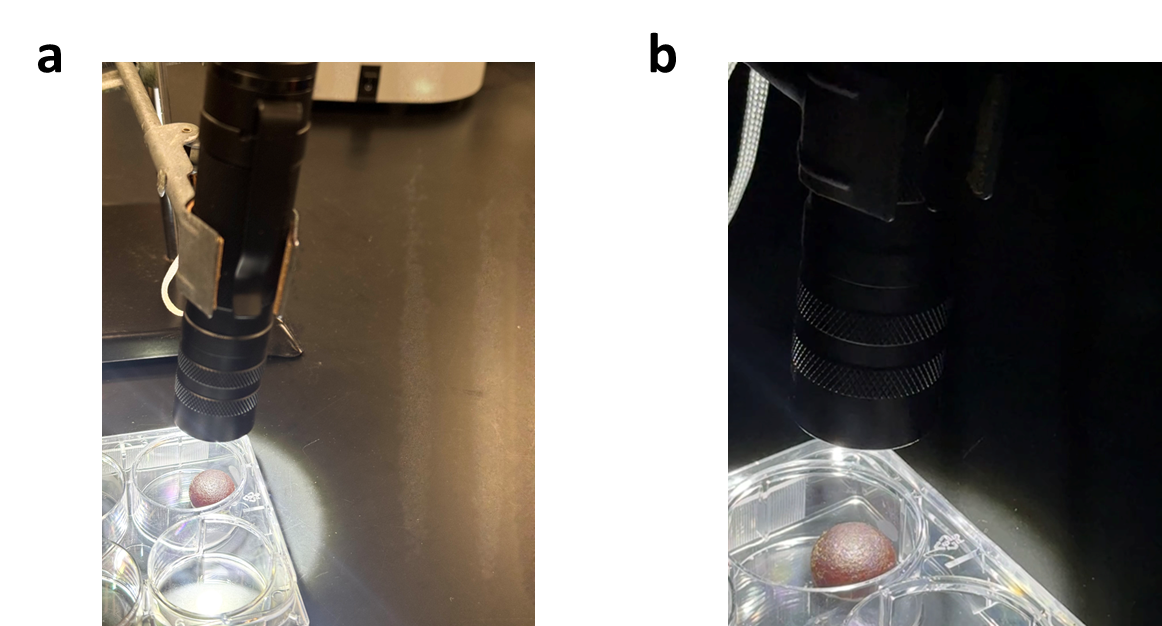

**Figure S27.** Experimental setup showing the photobleaching assessment study for PCA-PEG-ICG TMPs in (a) white light setting and (b) dark room.

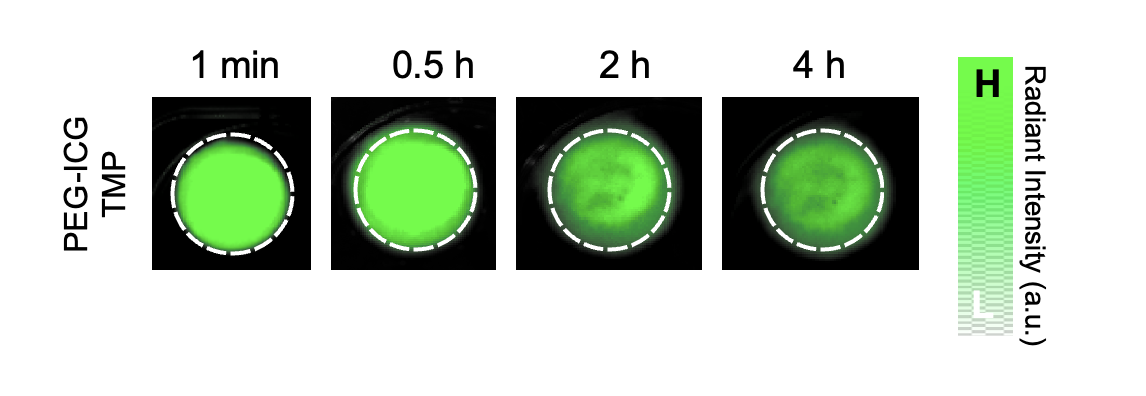

**Figure S28. NIR-I fluorescence imaging of PEG-ICG NP TMPs following white-light exposure.** Representative fluorescence images of PEG-ICG NP TMPs after exposure to a 1000-lumen white-light source for 1 min, 0.5 h, 2 h, and 4 h. Radiant intensity is shown in arbitrary units.

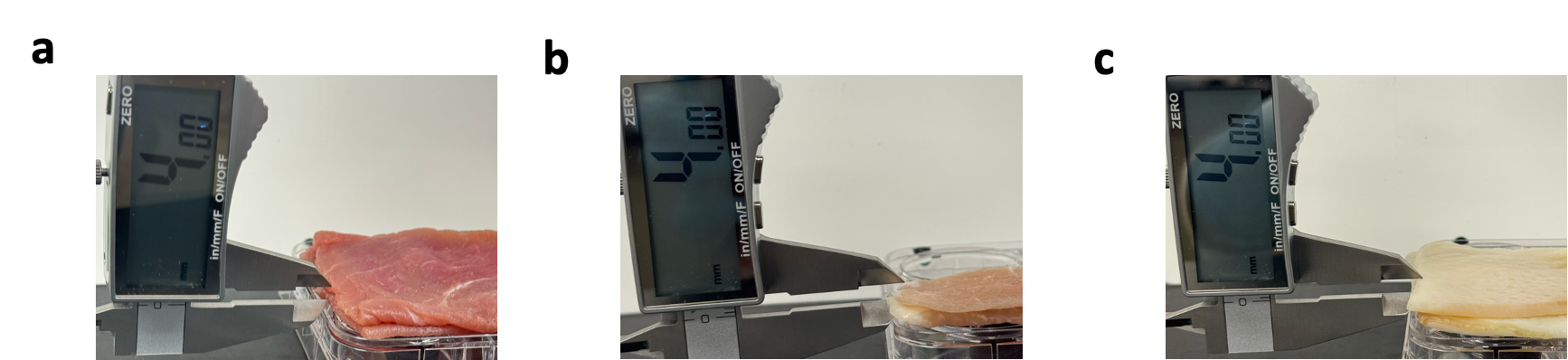

**Figure S29.** Representative images showing the stacked tissues: (a) muscle, (b) fat and (c) skin, respectively used for tissue stacking studies for PCA-PEG-ICG TMPs vs. ICG TMPs.

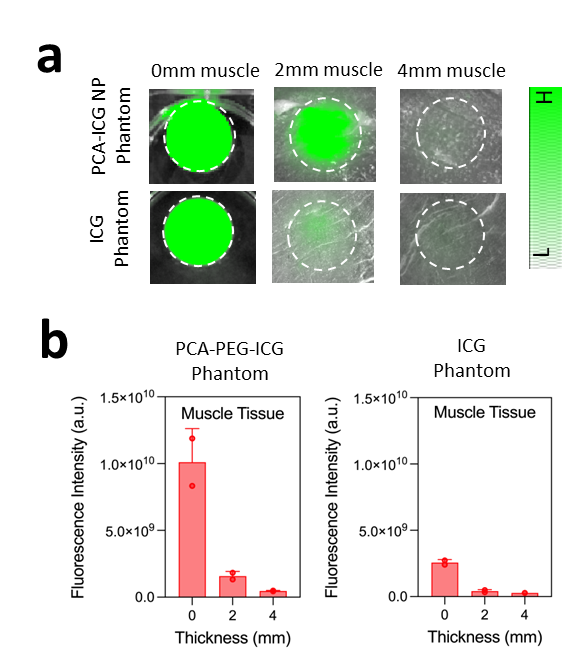

**Figure S30.** Tissue stacking profiling using muscle layers for TMPs. (a) Representative NIR-I images of PCA-PEG-ICG TMPs and free ICG TMP signal acquired under increasing muscle thickness (0, 2, 4 mm), illustrating signal attenuation with depth. (b) Quantitative analysis of radiant intensity demonstrates a progressive decrease in signal as tissue thickness increases. Data are presented as mean ± standard deviation (SD) (n=2).

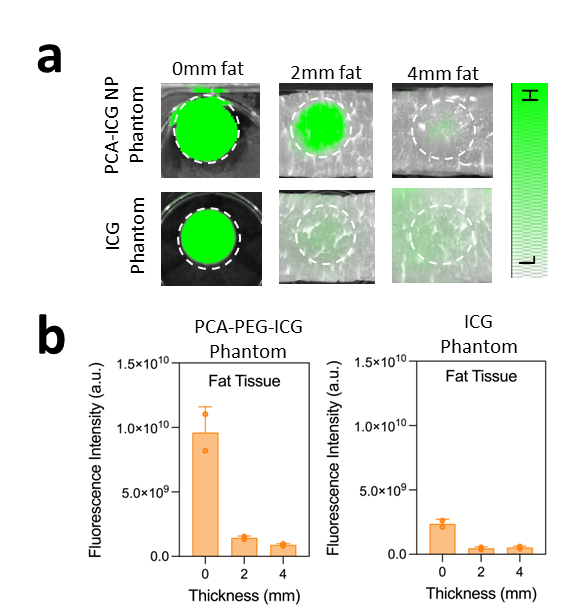

**Figure S31.** Tissue stacking profiling using fat layers for TMPs. (a) Representative NIR-I images of PCA-PEG-ICG TMPs and free ICG TMP signal acquired under increasing fat thickness (0, 2, 4 mm), illustrating signal attenuation with depth. (b) Quantitative analysis of radiant intensity demonstrates a progressive decrease in signal as tissue thickness increases. Data are presented as mean ± standard deviation (SD) (n=2).

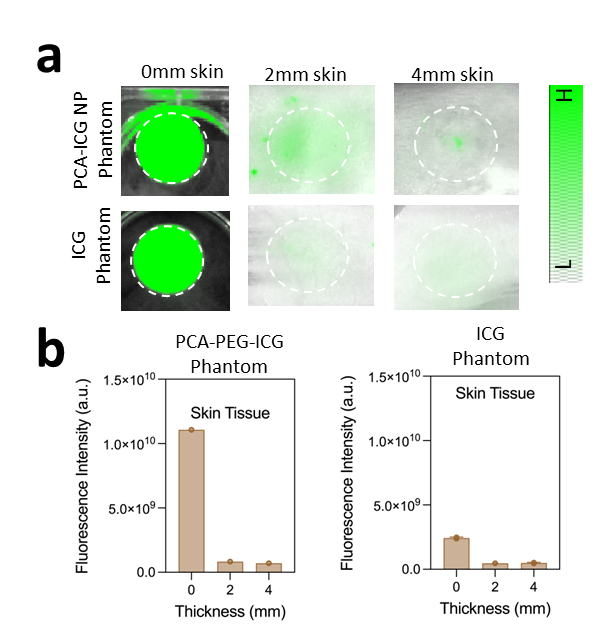

**Figure S32.** Tissue stacking profiling using skin layers for TMPs. (a) Representative NIR-I images of PCA-PEG-ICG TMPs and free ICG TMP signal acquired under increasing skin thickness (0, 2, 4 mm), illustrating signal attenuation with depth. (b) Quantitative analysis of radiant intensity demonstrates a progressive decrease in signal as tissue thickness increases. Data are presented as mean ± standard deviation (SD) (n=2).

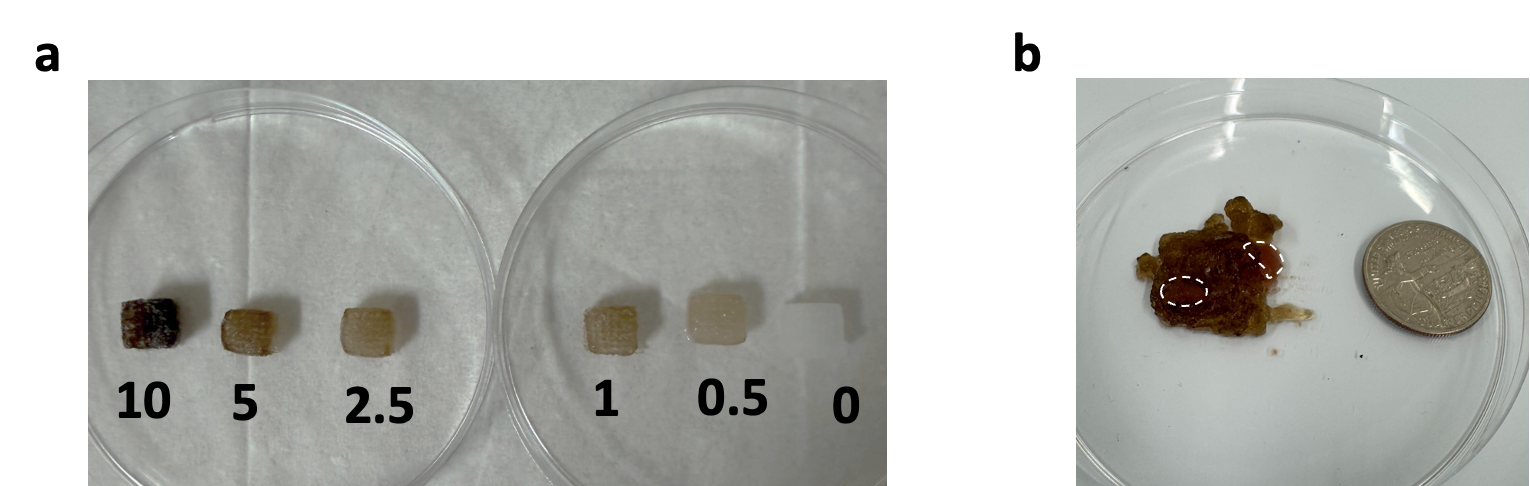

**Figure S33.** Skin mimicking constructs fabricated via 3D bioprinting with varying melanin concentrations (mg/mL), representing different levels of skin pigmentation.

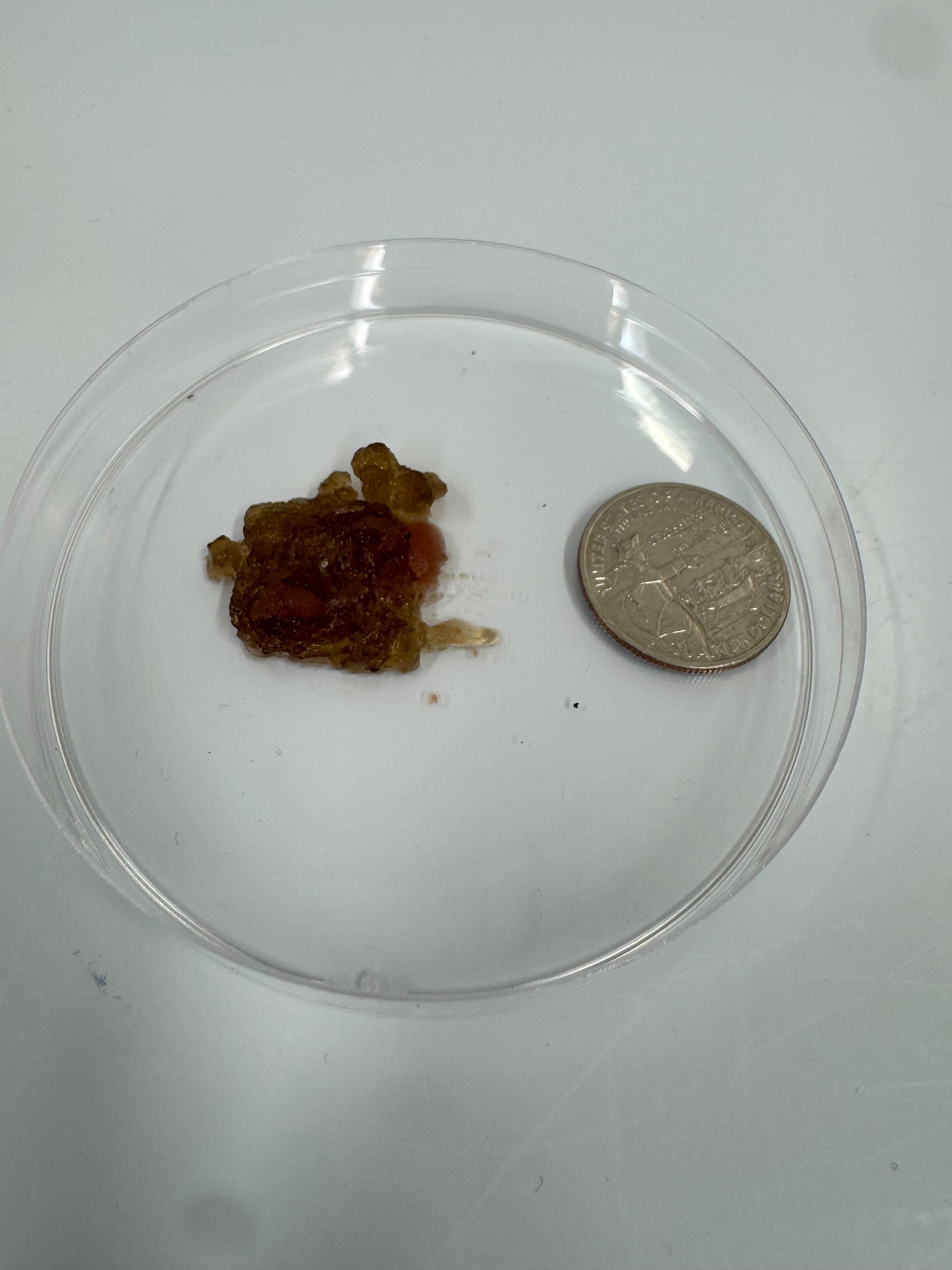

**Figure S34.** Image of 3D bio printed intraperitoneal (IP) tumor mimic consisting of multiple TMPs embedded on a skin-mimicking layer.
